# Fractional Amplitude of Low-Frequency Fluctuations Associated with Endocannabinoid, μ-Opioid and Dopamine Receptor Distributions in the Central Nervous System after High-Intensity Exercise Bouts

**DOI:** 10.1101/2023.10.06.561205

**Authors:** Henning Boecker, Angelika Maurer, Marcel Daamen, Luisa Bodensohn, Judith Werkhausen, Marvin Lohaus, Christian Manunzio, Ursula Manunzio, Alexander Radbruch, Ulrike Attenberger, Juergen Dukart, Neeraj Upadhyay

**Author notes:** <u>Correspondence to:</u> Univ.-Prof. Dr. med. Henning Boecker (https://orcid.org/0000-0003-2346-0598) Head of Clinical Functional Imaging Lab Department of Diagnostic and Interventional Radiology (Director: Prof. Ulrike Attenberger) University Hospital Bonn Venusberg Campus 1 53127 Bonn Germany.

## Abstract

Dopaminergic, opiod and endocannabinoid neurotransmission are thought to play an important role in the neurobiology of acute exercise and, in particular, in mediating positive affective responses and reward processes. Recent evidence indicates that changes in fractional amplitude of low-frequency fluctuations (zfALFF) in resting-state functional MRI (rs-fMRI) may reflect changes in specific neurotransmitter systems as tested by means of spatial correlation analyses. Here, we tested for this relationship at different exercise intensities in twenty young healthy trained athletes performing low-intensity (LIIE), high-intensity (HIIE) interval exercises and a control condition on three separate days. Positive And Negative Affect Schedule (PANAS) scores and rs-fMRI were acquired before and after each of the three experimental conditions. Respective zfALFF changes were analysed using a repeated measures ANOVAs. We explored spatial cross-correlations between pre-to-post zfALFF changes in each condition with available neurotransmitter maps using the JuSpace toolbox, and performed additional analyses for the main systems of interest (dopaminergic, opiod, endocannabinoid), focusing on specific brain networks related to ‘reward’ and ‘emotion’. Elevated PANAS Positive Affect was observed after LIIE and HIIE but not in the control condition. HIIE compared to the control condition resulted in differential zfALFF decreases in precuneus, orbitofrontal cortex, thalamus, and cerebellum, whereas differential zfALFF increases were identified in hypothalamus, pituitary, and periaqueductal gray. The spatial alteration patterns in zfALFF were positively associated with dopaminergic and μ-opioidergic receptor distributions within the ‘reward’ network. These findings provide new insight into the neurobiology of exercise supporting the importance of reward-related neurotransmission during high-intensity physical activity.

**Keypoints:**

1. Positive mood changes, indexed as elevated PANAS Positive Affect, were identified after high- and low-intensity exercise bouts, supporting previous accounts on mood-improving effects of physical activity.
2. High-intensity exercise was found to be associated with distributed changes in fractional amplitude of low-frequency fluctuations, indicating enduring neural activity changes after anaerobic exercise bouts.
3. Results of spatial cross-correlations with representative PET neurotransmitter distribution maps suggest involvement of endocannabinoid, dopaminergic, and opioidergic neurotransmission after high-intensity exercise.
4. Utilizing spatial cross-correlations of changes in fractional amplitude of low-frequency fluctuations and representative PET neurotransmitter distribution maps, despite being an indirect metric, provides an innovative methodological framework for human exercise research, as it allows for non-invasive testing of acute exercise-related changes multiple neurotransmitter.

## INTRODUCTION

Physical activity is considered one of the most efficient mood-regulating behaviors [1]. Much work has been undertaken to unravel the underlying mechanisms of affective changes triggered by physical activity in animal models and in a growing number of human studies [2]. Various studies show that moderate exercise regimens induce acute improvements of positive affect, especially immediately after exercise [3]. Moreover, there is some evidence that acute exercise can exert motivational effects [4], including phenomena of transient appetite suppression or altered reactivity to food cues [5–7].

Meanwhile, the underlying changes in regional brain activity remain understudied. Considering cerebral blood flow (CBF) as an important surrogate marker for brain activity, human evidence mainly comes from transcranial doppler sonography or near infrared spectroscopy studies, and suggests at least transient perfusion increases which may plateau or return to baseline, depending on duration and intensity of exercise bouts, among other factors [8]. But these methods provide no (or limited) information about region-specific effects, and there are theoretical accounts which assume that resource limitations may also necessitate temporary downregulation of certain brain regions (e.g. the reticular-activating hypofrontality model: [9]). Indeed, preliminary evidence from magnetic resonance imaging (MRI) studies using arterial spin labeling (ASL) indicates post-exercise CBF increases in young adults for the hippocampus [10] or posterior insula, but also concomitant CBF decreases in the medial orbitofrontal cortex and dorsal striatum [6]. Indirect evidence for region-specific activity changes also comes from resting-state functional MRI (rs-fMRI) studies. Using blood oxygenation-dependent (BOLD) fMRI, most studies examined post-exercise changes in functional connectivity (FC), i.e. the spatiotemporal coherence of spontaneous brain activity fluctuations between different brain areas which are interpreted to reflect integrated functional brain networks [11]: This includes exercise-related increases FC in sensorimotor networks [12], or affect-reward, hippocampal, cingulo-opercular, and executive control networks [13], but also contrary findings of FC reductions [14]. Exercise intensity may also play a moderating role [11], as suggested by studies observing differential pattern of FC increases and decreases in specific rs-fMRI networks after low- or high-intensity exercise bouts [15, 16]. Still, rs-fMRI can provide additional brain activity-related markers, including the amplitude of low-frequency fluctuations (ALFF). It represents the voxel-level magnitude of regional BOLD fluctuations in the low-frequency range (0.01–0.08 Hz) which is assumed to be proportional to neural activity [17]. Fractional ALFF (fALFF) is a normalized index of ALFF which is considered to be less sensitive to physiological noise [18]. Previous data showed a close association with underlying metabolic activity [19], making it a possible surrogate marker for changes in regional brain activity. While one recent study [20] examined the cross-sectional relationship between fALFF measures and cardiorespiratory fitness measures in trained athletes, complementary applications in acute exercise studies are missing.

Neurotransmitters and neuromodulators are thought to play a crucial role in mediating the mood-regulating effects of exercise, and changes in affective homeostasis, anxiety and depression have been commonly associated with monoamine [21], endorphin [22], and endocannabinoid [23] neurotransmission. In addition, the motivation to exercise and to overcome physical boundaries and pain, as is particularly the case for strenuous high-intensity and/or long-duration exercise regimens, has been particularly linked to the mesolimbic dopamine reward circuit [24, 25]. As yet, most of the current knowledge on exercise-induced central neurotransmission has been accumulated in animal studies [2] using either *ex vivo* brain tissue assays with high-performance liquid chromatography/mass spectrometry [26, 27], Western blot and immunofluorescence [28], and autoradiography [29]; or *in vivo* sampling with microdialysis [27, 30, 31]. In humans, *in-vivo* examinations have been conducted based on selective pharmacological manipulations to clarify the roles of dopaminergic [32–36], opioidergic [37, 38], and endocannabinoid [38] neurotransmission in mediating exercise-induced affect modulation and reward-related behavior.

Furthermore, the last 20 years have seen the introduction of functional neuroimaging into the field of exercise science, with positron emission tomography (PET) holding unique potential for localizing and quantifying training-induced neurotransmission *in vivo* after acute training sessions, or long-term changes in receptor distribution after repetitive training [39, 40]. PET ligand displacement studies allow *in vivo* monitoring of endogenous transmitter trafficking at the human whole-brain level after acute exercise bouts and, thereby, to identify the link between exercise-induced behavioral measures and endogenous neurotransmission [41]. Focusing on the dopaminergic system, Ouchi et al. [42] demonstrated that a gait challenge significantly increased endogenous dopaminergic release. A more recent PET ligand displacement study with [11C]Raclopride reported that dopamine release after steady-state cycling was more pronounced in the ventral striatum in habitual exercisers than in sedentary subjects [43], suggesting an influence of training status. The opioidergic system has been studied using a nonselective opioid ligand [22]. After 2 h of endurance running, ligand displacement effects were described in prefrontal and limbic/paralimbic brain structures which were also inversely correlated with the level of post-exercise euphoria, hence, suggesting a link between opioid release in affect-related brain areas and affective states after physical exercise [22]. Subsequent studies have confirmed increased opioidergic neurotransmission after acute exercise, particularly central μ-opioidergic activation using [11C]carfentanil PET, with evidence for exercise intensity-dependent effects [44, 45]. Moreover, [11C]Carfentanil PET indicated aerobic exercise to modulate anticipatory reward processing via the μ-opioid receptor system [46]. Endocannabinoids like anandamide (AEA) and 2-arachidonoglycerol (2-AG) are endogenous ligands for cannabinoid (CB1) receptors that are densely expressed in cortico-limbic brain networks involved in emotional control, such as amygdala, hippocampus and cortical areas [47]. Moreover, there are meta-analytical indications that endocannabinoids play a causal role for improvements in mood and affect following aerobic exercise [48, 49]. It has been shown that exercise increases circulating endocannabinoid levels in an intensity-dependent manner [50, 51]. One PET study in humans employed the CB1 inverse agonist radioligand [18F]FMPEP-d_2_ in individuals with risk for obesity, indicating links between reduced CB1 availability and poor exercise habits [52]. However, to the best of our knowledge, no PET endocannabinoid tracer studies on acute or long-term exercise challenges with a pre/post design have been published so far.

It is important to point out that despite the genuine interest in PET displacement studies during exercise challenges, studies are hampered by limited availability, cost, and radiation exposure [39–41]. The latter aspect is the main limiting factor preventing a comprehensive analysis of the effects of exercise on several transmitter systems at the same time. Indeed, pre-post designs are largely limited to one or maximally two neurotransmitter systems due to radiation exposure limitations, especially in healthy young subjects. Recent rs-fMRI studies have shown that spatial fALFF patterns are associated with the distribution of specific receptor systems targeted by respective PET and SPECT compounds [53] and that this approach cannot only be used to make inferences about the status of various neurotransmitter systems (e.g., in neurological disorders), but also to detect acute changes in neurotransmission in relation to pharmacological challenges. In an extension of this latter rationale, the current study intended to investigate how varying intensities of acute exercise affect fALFF changes in brain regions linked to representative PET and SPECT neurotransmitter maps [53, 54], thereby providing indirect markers for their involvement in acute exercise. Following up on previous work suggesting intensity-dependent effects on rs-fMRI measures [15, 16], the current study examined fALFF map changes as means to investigate associated neurotransmitter effects in trained male athletes after performing a control, low-intensity (LIIE), and high-intensity (HIIE) interval exercise session. Complementing a traditional voxel-wise analysis to identify individual brain regions showing exercise-related differences in brain activity (as indicated by fALFF), we tested the spatial correlation between the observed changes across regions and the distribution of neurotransmitter systems to make inferences about the exercise-induced activation of the latter. We hypothesized specific involvement of dopaminergic, μ-opioidergic, and endocannabinoid neurotransmission, in particular for the high-intensity condition [44, 45, 50]. Meanwhile, we also examined potential relationships with other neurotransmitter systems provided by the JuSpace toolbox [53] in an exploratory manner. Considering previous human exercise studies which indicated exercise-induced effects in reward- [4, 46] and affect- [22, 55] related brain networks, we also performed separate analyses focusing on those regions which overlap with meta-analytically defined ‘emotion’ and ‘reward’ networks.

## METHODS

### General Study Design

This study, referred to as BEACON (‘**B**icycling **E**ffects on **A**ffect and **CO**gnition in **N**euroscience’), was conceptualized as a within-subject design to investigate the acute effects of exercise bouts of differing exercise intensities on brain functional networks as well as affect and cognition (**Figure 1**). Some of the reported measures will also be included in other manuscripts derived from this study with different research questions.

**Figure 1:**
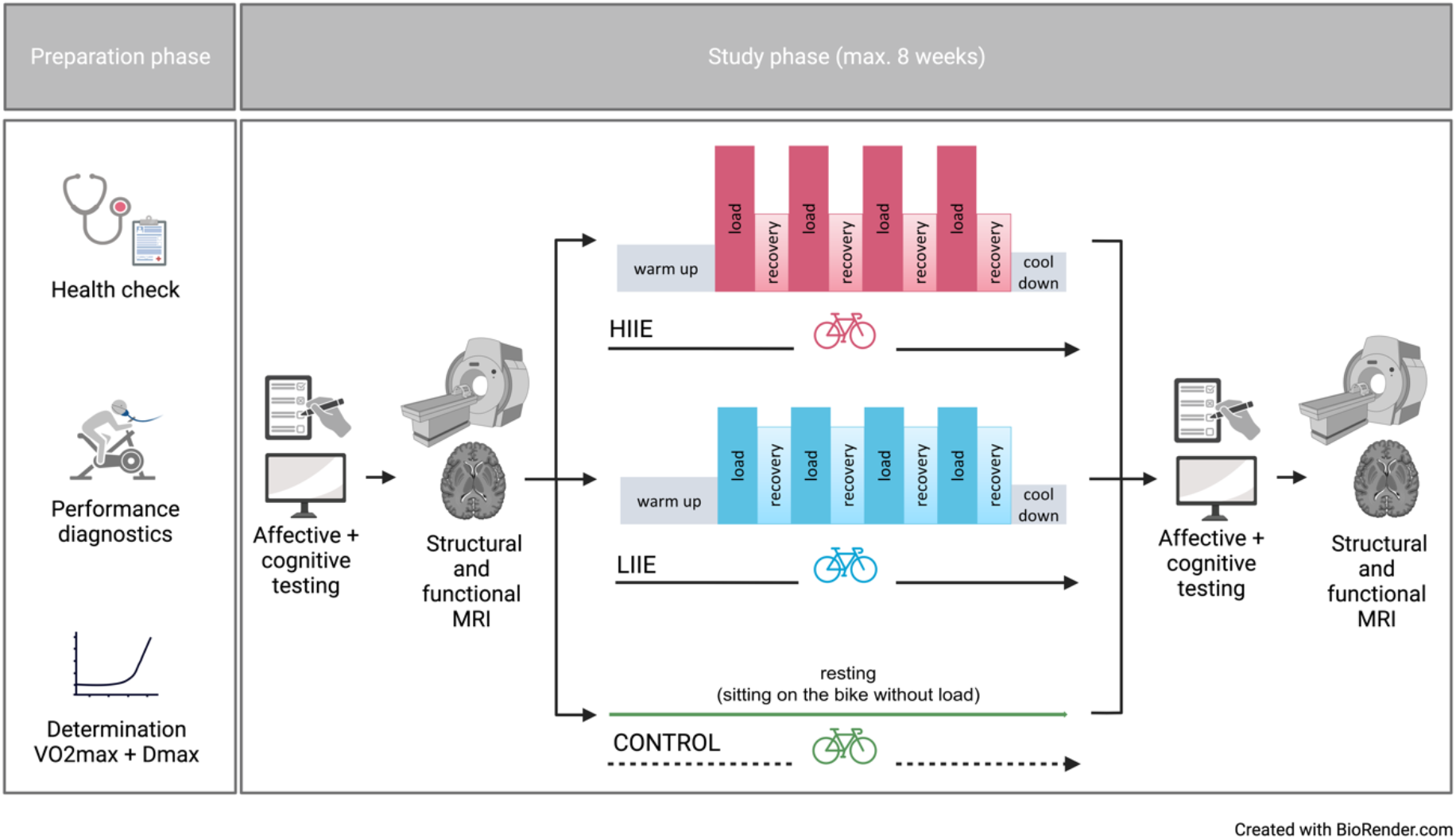
General Study Design.

All participants were informed about the study protocol, examinations, potential discomforts, risks and a written informed consent was obtained. Ethical approval was given by Ethics Committee at the Medical Faculty of the Rheinische Friedrich-Wilhelms-Universität Bonn (Nr. 358/19), conferring to national legislation and the Declaration of Helsinki.

Each participant completed three distinct intervention conditions - aerobic cycling (low intensity; LIIE), anaerobic cycling (high intensity; HIIE), and rest (control) condition - in a randomized order (further information below). The interventions were carried out within a maximum of 8 weeks and at least 7 days apart. The night before the test, the subjects were instructed to obtain enough sleep, refrain from drinking alcohol for the preceding 24 hours, and hold off on caffeine until two hours before the test. Additionally, subjects were told not to exercise 24 hours beforehand. To prevent measurement differences owing to anxiety on the first exam day, subjects experienced a mock scanner session before the actual MRI. Other than that, the processes were carried out in a uniform manner on each examination day. First, subjects completed general and mood-related questionnaires as well as cognitive tests (a condensed version of the Attention Network Test (ANT) [56] and object recognition memory performance/hippocampal integrity using the Mnemonic Similarity Task (MST) [57] were gathered, but were not subject of the current analyses). Then, rsfMRI was acquired. Subsequently, one of the three interventions was performed. After the intervention, mood questionnaires, cognitive tests as well as fMRI were repeated in the same order as pre-intervention.

### Participants

Well-trained right-handed male athletes aged between 20-35 years were recruited via flyer distribution at local cycling and triathlon clubs as well as the university hospital and social media. Subject enrollment was restricted to male participants in order to minimize variance due to hormonal fluctuations during the menstrual cycle. Individuals with current and/or previous severe psychiatric, neurologic or cardiovascular diseases in addition to the typical MRI-specific exclusion criteria i.e., claustrophobia, non-removable metal/implants, tattoos exceeding a critical size or other prohibiting reasons were excluded. Additionally, in order to ensure a high fitness level of participants, individuals with a relative maximum oxygen uptake (relVO_2max_) below 55 ml/min/kg were excluded [58].

### Experimental Procedures Preparation Phase

#### General Screening

Selection was performed prior to the study phase: It included the recording of sociodemographic characteristics, a sports medical examination and performance diagnostics. A sociodemographic questionnaire and vocabulary test were administered to obtain descriptive characteristics such as age, educational level, and estimated verbal intelligence. The International Physical Activity Questionnaire (IPAQ) [59] was carried out to assess the physical activities of participants in everyday life. Edinburgh-Handedness-Inventory [60] was used to assess handedness. Further, participants were screened for psychiatric symptoms using several questionnaires such as the Mini International Neuropsychiatric Interview [61], the Beck Depression Inventory (BDI) [62], and the trait anxiety of the State-Trait-Anxiety Inventory (STAI) [63]. Moreover, a substance abuse questionnaire excluded possible substance use disorders.

The medical sports examination included an anamnestic questionnaire, lung and cardiac auscultation, and a 12-lead resting ECG. Where necessary, an additional transthoracic echocardiogram was carried out.

#### Performance Diagnostics

Performance diagnostics, which included a maximal incremental step test on a cycling ergometer (Cyclus2, RBM Elektronik-Automation GmbH, Leipzig, Germany), were carried out to ascertain physical fitness and, consequently, individual training intensities. The Cyclus2 made it possible to mount each participant’s own bike, guaranteeing that each subject would cycle in the same unique position throughout the study. The initial workload for the incremental step test was 100 watts (W), followed by 20 W increases every three minutes (min) at a cadence of 80 revolutions per minute (rpm) until volitional exhaustion. Power was digitally controlled with direct drive throughout the performance diagnostics and the ensuing exercise modifications. When the cadence dropped to less than 65 rpm, the test was stopped.

Throughout the test, measurements of oxygen uptake (VO_2_), the metabolic respiratory quotient (RER), heart rate (HR) (Cortex meta-analyzer 3B, Leipzig, Germany, and Polar Electro Oy, Kempele, Finland), and the electrocardiogram (ECG) (Cardio 100 USB, Ergoline GmbH, Bitz, Germany) were continually recorded. A validated H10 chest strap was used to continually measure HR variability. Additionally, blood pressure was checked at each step. The final 15 seconds of each procedure were used to draw 20 µl of capillary blood from the earlobe, mix it immediately with 1 mL of the EBIO plus system hemolysis solution, and analyze the sample amperometrically and enzymatically using EBIOplus (EKF Diagnostic Sales, Magdeburg, Germany). Within the last 15 seconds of each increment, the rating of perceived exertion (RPE) was measured using the 6 to 20 point Borg scale (6 no exertion at all, 20 maximal exertion) [64].

Exhaustion was considered when at least two of the following criteria were met: plateauing in VO_2_, RER ≥ 1.05, high levels of blood lactate (≥8 mmol/L), a RPE of ≥18. The greatest 30-second moving average of VO_2_ divided by body mass (mL/min/kg) was used to calculate relVO_2max_. Those who had a relVO_2max_ of less than 55 mL/min/kg were not included in the study.

### Study Phase

#### Intervention

Following the first MRI examination, volunteers completed the supervised intervention on a bicycle ergometer. Depending on the treatment condition, subjects performed one of the two alternative exercise intensities (low and high) or the control condition (without load) in randomized order. An adapted Latin Square was used for randomization: ABC, ACB, BCA, BAC, CBA, CAB; order of these combinations additionally randomized within blocks of 6 subjects. A 4*4-minute load was alternated with 3 minutes of active recovery during the exercise interventions. Individual exercise intensities were established using the incremental step test. In the exercise interventions, subjects began with a 10-minute warm-up at 1,5 watts per kilogram of body weight (W / kg BW) followed by interval training and a final cooldown of 5 minutes <1,5 W / kg BW. The intervention intensities were established as follows: i) LIIE: 4 * 4 minutes’ load at 100% first rise (W) with 3 minutes active recovery at 90% of first rise; ii) HIIE: 4 * 4 minutes’ load at 110% D_max_ (W) with 3 minutes active recovery at 60% of D_max_, and iii) control: no load while sitting on the cycling ergometer for 43 minutes.

The modified D_max_ approach of Zwingmann et al. [65] was used to calculate lactate thresholds. The first rise was defined as the moment at which the lactate concentration increased by >4% from the previous value. The last 30 seconds of each load and recovery period were used to measure blood lactate, HR_int_ (during the intervention), and RPE. Blood pressure was tested both before and after the treatment.

#### Physiological and Psychological Testing

Each study day began with the subjects filling out a special questionnaire that measured their current levels of exhaustion, discomfort, previous night’s sleep quantity and quality, and recent caffeine and alcohol intake.

To account for physiological effects due to exhaustion, the corresponding MoodMeter [66, 67] subscale (consisting of the items “feeble” and “drowsy”) from the psychological strain was included in the statistical analyses as this may influence mood and fMRI outcomes.

The Positive And Negative Affect Schedule (PANAS) was one of the questionnaires used to gauge mood in general [68], which served as the primary affect measure in the presented analysis. Participants had to rate their current mood state based on a total of 20 adjectives, 10 of which are assigned to the dimension of positive affect and 10 to the dimension of negative affect (scoring with 5-point Likert scale ranging from 1 ‘not at all’ to 5 ‘very much’). This way, the questionnaire allowed for a separate dimensional assessment of pleasure and displeasure effects which are assumed to be partially dissociable on the brain system level [69], and may therefore show differential susceptibility to acute exercise. To examine potential anxiolytic effects of exercise more specifically, the STAI-State scale was also collected [63].

#### MRI Acquisition

On each study day, participants completed two identical MR sessions. The respective MRI scans (in total six scans) were acquired on a Philips Ingenia Elition 3.0T with a 32-channel head coil at the Parent-Child Centre of the University Hospital Bonn.

A regular session consisted of a T1-weighted (T1w), a fieldmap and a rs-fMRI sequence. For rs-fMRI, the light in the examination room was switched off and participants were instructed to close their eyes, not to think of anything in particular, and to stay awake during the whole scan. An echo-planar imaging (EPI) protocol with blood-oxygen-level-depending (BOLD) contrast and 3D acquisition was performed with the following specifications: TR = 1020ms, TE = 30ms, acquired voxel size = 2.5×2.5×2.5mm, reconstructed voxel size = 2.17 x 2.17 x 2.5 mm, FoV = 208×208mm, flip angle = 52°, SENSE: 2, MB factor: 3, EPI factor: 41, matrix: 84 x 82, slices: 51, scan order: FH (ascending). Over a total duration of 10:04 min, 585 dynamic scans were acquired. The anatomical T1w sequences were acquired with the following specifications: TR = 10 ms, TE = 4.7 ms, acquired voxel size = 0.7×0.7×0.7mm, reconstructed voxel size: 0.49×0.49×0.57 mm, FoV = 250×250mm, flip angle = 8°. The total duration of this sequence was 6:19 min. Field mapping was performed with the following parameters: TR = 650 ms, TE = 7 ms, voxel size: 3.75 x 3.75 x 4 mm, flip angle 80°.

During all fMRI scans, HR was continuously recorded (HR_rest_) by a pulsoxymetry sensor on the index finger.

### Physiological and Psychological Data Analysis

The analysis of the behavioral and the physiological data was performed using IBM SPSS (Statistics Version 27.0. IBM Corp. Armonk, NY).

#### Exercise Intervention

To evaluate whether exercise intensity between the low and high condition differed significantly, paired t-tests with Bonferroni correction were performed for the variables HR_int_, lactate concentration and RPE. Significance was considered at p<0.05 and the effect size is reported as Cohen’s d.

#### HR_rest_ and Exhaustion

Values above or below the mean ± 2.5 standard deviations (SD) were removed from the data set as outliers for HR_rest_. Statistical analysis of the outlier-corrected means of HR_rest_ and raw exhaustion scores was performed using a 2 (timepoint: pre/post) x 3 (condition: control, low, high) repeated measures ANOVA, separately. Post-hoc paired t-tests with Bonferroni correction were also performed. Significance was considered at p<0.008 (0.05/6), and the effect size was reported as partial eta square and Cohen’s d.

#### Mood Questionnaires

A repeated measures 2 (timepoint: pre-intervention/post-intervention) x 3 (condition: control/ low/ high) ANOVA was performed to assess the changes in PANAS and STAI-State according to the conditions. Post-hoc paired t-tests with Bonferroni correction were also performed. Significance was considered at p<0.008 (0.05/6), and the effect size was reported as partial eta square and Cohen’s d.

### MRI Data Analysis

#### Qualitiy Control and Preprocessing

Resting state fMRI data underwent a well-recognized MRIQC [70] pipeline for visual quality check regarding acquisition artifacts.

Resting state fMRI data were pre-processed using the fmriprep pipeline (https://fmriprep.org/en/stable/) [71]. Preprocessing started with skull stripping the fMRI data using a custom fmriprep methodology. Furthermore, fieldmaps were used to correct for susceptibility distortions by applying the FSL 6.0 [72] fugue (https://fsl.fmrib.ox.ac.uk/fsl/fslwiki/FUGUE) and SDCflows tools [73]. The estimated susceptibility distortion was used to create a corrected BOLD EPI reference in order to accurately register the functional data to the anatomical reference. The BOLD EPI reference was registered to the T1-weighted reference using a boundary-based registration approach. Before spatiotemporal filtering, head motion parameters with respect to BOLD EPI reference were estimated using mcflirt [74]. A single composite transform was applied to the BOLD time-series to correct for head motion and susceptibility distortions, and resampled to their original native space. Thereafter, BOLD time series were normalized to standard MNI152NLin2009cAsym space [75]. A detailed description of the fmriprep pipeline can be found in the Supplement (‘Detailed Description of the fMRIPrep Pipeline (Boilerplate)’).

After preprocessing, further data quality was determined on the basis of the fmriprep DVARS and framewise displacement (FD) metrices. Both DVARS and FD reflect the rate of change of the BOLD signal across the whole brain at each frame of data and the head movement of the specific frames, respectively. In the case of more than 60% of the 585 volumes with FD larger than 0.2 mm, subjects were removed from the final analysis [76].

Then, output functional maps were used to calculate fALFF for all six sessions (pre- and post-intervention in each condition). The 3dRSFC function in AFNI (https://afni.nimh.nih.gov/pub/dist/doc/program_help/3dRSFC.html) was applied, including quadratic detrending, band-pass filtering (0.01–0.08 Hz), 4mm smoothing and regressing out white matter and cerebrospinal fluid signal as well as the 24 motion parameter time courses in a single step (Taylor and Saad, 2013). Normalized fALFF (zfALFF) maps were created by dividing the map through its mean.

#### Voxel-wise Analyses of zfALFF Maps

In a basic analysis, we compared the zfALFF maps from the pre- and post sessions of the three conditions using a voxel-wise approach performing a 2 (timepoint: pre/post) x 3 (condition: control, low, high) repeated measures analysis of variance (ANOVA) for whole brain in SPM12 (https://www.fil.ion.ucl.ac.uk/spm/software/spm12/).

To exclude a direct effect of HR_Rest_ and Exhaution on the zfALFF, regression analyses were performed with the zfALFF maps within each condition (HIIE, LIIE, and control) for the post-pre changes. Masks were created by saving significant clusters found at a lenient threshold of p<0.01 (uncorrected). These masks from each condition were merged to one mask indicating brain regions influences by HR_Rest_ and Exhaution. Later, the merged mask was used for exclusive masking in the repeated measure ANOVA for zfALFF.

We applied a voxel-wise cluster-defining threshold of *p* < 0.001 (uncorrected) and reported significant clusters at *p* < 0.05 after implementing a cluster-wise threshold for different contrasts using the family-wise error (FWE) method for multiple comparisons correction.

#### Spatial Correlations of fALFF changes with Neurotransmitter Maps

The JuSpace toolbox (https://github.com/juryxy/JuSpace; version 1.4) was used to compute spatial cross correlations between the zfALFF changes per exercise condition (within-subject pairwise zfALFF differences between the pre- and post-intervention fMRI scan; computing option 6 of the toolbox) and neurotransmitter maps.

First, we tested for spatial correlations of the zfALFF changes induced by either of the three conditions (LIIE, HIIE or control) with different neurotransmitter systems using a whole brain approach, using 14 neurotransmitter maps included in the JuSpace toolbox. In case of multiple maps per neurotransmitter system, the ones with highest reliability based on sample size and/or signal-to-noise ratio were chosen. We included the following neurotransmitter maps: dopamine synthesis, storage and transport (Fluorodopa [77]; DAT [78]) and receptors (D1 [79]; D2 [54, 80], GABA (gamma-aminobutyric acid) receptor [78], endocannabinoid receptor (CB1) [54, 81], μ-opioidergic receptor (MOR) [54, 82], serotonin receptors (5-hydroxytryptamine 1a (5-HT1a, 5-HT1b, and 5-HT2a) and transporter (SERT) [83], noradrenaline transporter (NET) [84], vesicular acetyl choline transporter (VAChT) [54, 85], and the metabotropic glutamate receptor type 5 (mGluR_5_) [54, 86].

Second, we tested for spatial correlations of the zfALFF changes in those regions of the neurotransmitter maps which overlapped with *a priori* defined ‘reward’ and ‘emotion’ networks. For this hypothesis-driven approach, we focused on three pre-informed neurotransmitter systems (i.e. D2, Mu and CB1) already known to be involved in affect and reward modulation in general [87, 88], or in the exercise context more specifically [22, 46, 48, 52, 89]. For these analyses, spatial correlations were restricted to brain regions within the neurotransmitter-specific maps which are also linked with emotion and reward processing, respectively, based on independent neuroscientific evidence: A ‘reward network’ mask was created using a term-based meta-analysis of functional neuroimaging studies implemented in Neurosynth (https://neurosynth.org) including the ventral striatum, thalamus, cingulate, orbitofrontal cortex. An ‘emotion network’ mask was applied in an identical manner as in our previous study [16] and included anterior/middle/posterior cingulate cortex, inferior/medial/middle/superior orbitofrontal cortex, dorsolateral prefrontal cortex, hypothalamus, insula, amygdala, nucleus accumbens, and pallidum. Both masks are displayed in the Supplement (**Figure S1**).

Spearman correlation coefficients were calculated between the single subject pairwise zfALFF difference maps and the respective neurotransmitter maps, after parcellation according to the Neuromorphometric atlas [90].

Exact permutation-based p-values (with 10.000 permutations) were computed for all comparisons to test for significance of the mean correlation coefficients observed across participants from the null distribution. Finally, we reported the spatial cross correlations at p < 0.05, false discovery rate (FDR) corrected for multiple comparisons.

#### fALFF-neurotransmitters Correlations with Mood Questionnaires

Finally, we used Spearman correlations to assess the relationships between the outputs of JuSpace (significant zfALFF-neurotransmitters correlation, Fisher’s Z-transformed) and the PANAS and the STAI-State, respectively. Significant results were reported at p<0.05 after correcting for multiple comparisons using Bonferroni correction.

## RESULTS

### Demographic Measures

Among twenty-nine recruited participants only twenty (age: 27.3 ± 3.55 years) were included in the present analyses. Reasons for exclusion were: injury during private exercise or illness (N=4), failure to reach the specified VO_2max_ limit of 55 mL/min/kg (N=3), and motion artefact in MRI (N=2). An overview of demographic and physiological characteristics of the sample is provided in **Table 1** and in other manuscripts from this work. According to the IPAQ, N=19 subjects could be categorized as highly active and N=1 as moderately active.

**Table 1:**
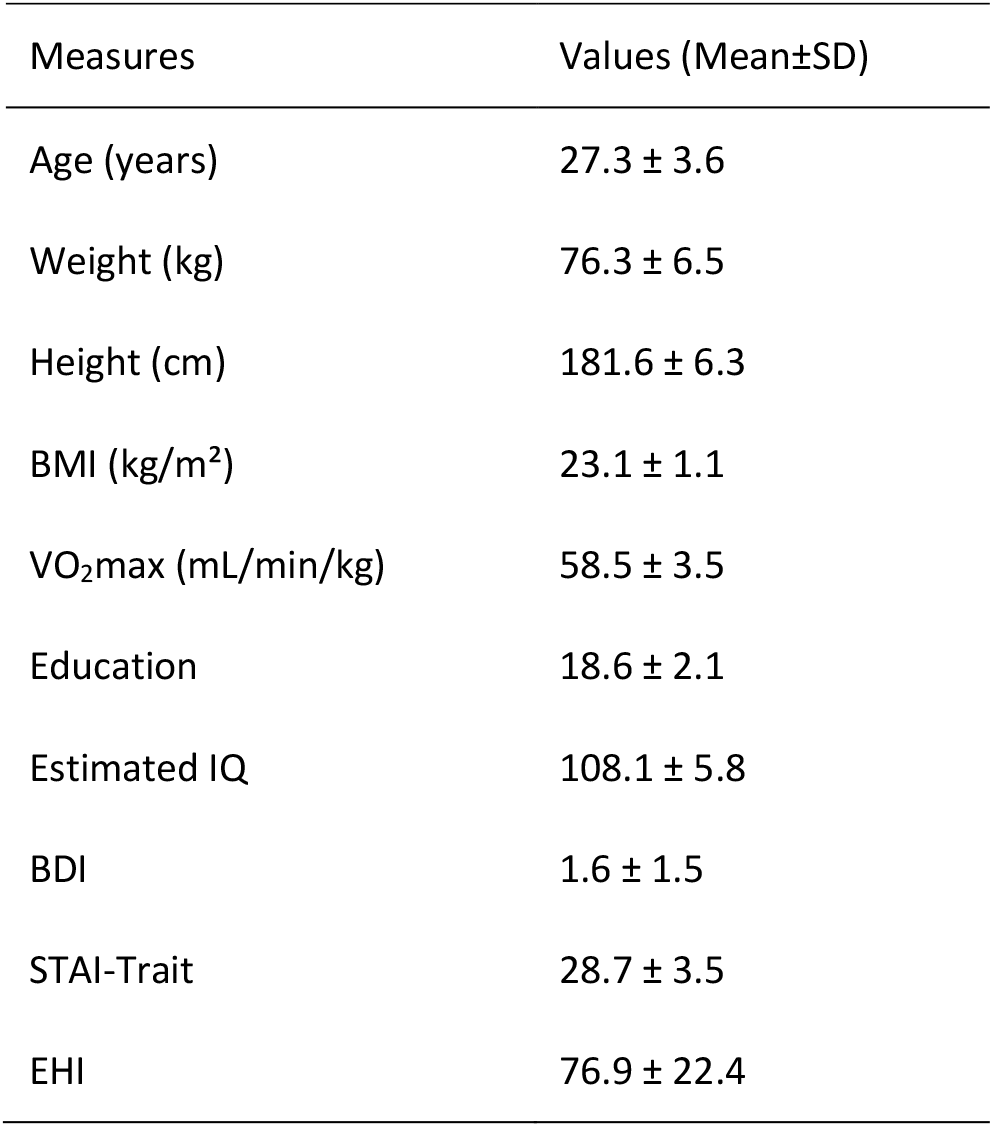
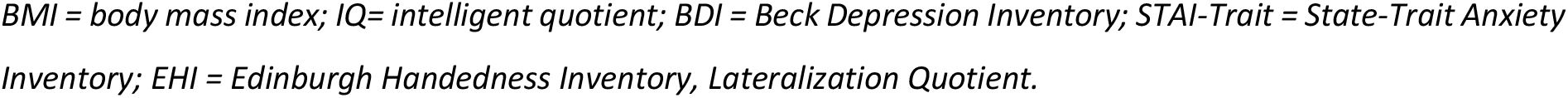
Demographic and physiological characteristics.

| Measures | Values (Mean±SD) |
| --- | --- |
| Age (years) | $27.3 \pm 3.6$ |
| Weight (kg) | $76.3 \pm 6.5$ |
| Height (cm) | $181.6 \pm 6.3$ |
| BMI (kg/m<sup>2</sup>) | $23.1 \pm 1.1$ |
| $VO_{2max}$ (mL/min/kg) | $58.5 \pm 3.5$ |
| Education | $18.6 \pm 2.1$ |
| Estimated IQ | $108.1 \pm 5.8$ |
| BDI | $1.6 \pm 1.5$ |
| STAI-Trait | $28.7 \pm 3.5$ |
| EHI | $76.9 \pm 22.4$ |

*BMI = body mass index; IQ= intelligent quotient; BDI = Beck Depression Inventory; STAI-Trait = State-Trait Anxiety Inventory; EHI = Edinburgh Handedness Inventory, Lateralization Quotient.*

### Physiological Data

Results of the HR_int_ and HR_rest_ are also reported in other manuscripts from this study (in preparation / parallel submission) with different research questions. Since the intervention is a key component of the study, overlapping reporting is inevitable.

#### Exercise intervention

All included participants were able to train within the desired power output in the LIIE and the HIIE interventions. Statistical analysis revealed significant differences between the two exercise conditions. Values and statistics of HR_int_ (**Table S1**), lactate concentration (**Table S2**), RPE (**Table S3**), and the actual power values during the intervention (**Table S4**) are summarized in the Supplementary Materials.

#### Heart rate during fMRI (HR_rest_)

The results of the repeated measures ANOVA following Greenhouse-Geisser method for sphericity correction showed a significant main effect of condition (F(2, 32.36) = 6.80, p = 0.005, η^2^= 0.26), a main effect of time (F(1, 19) = 7.02, p = 0.016, η^2^= 0.27), and a significant time × condition interaction (F(2, 26.06) = 18.01, p < 0.001, η^2^= 0.49) for the variable HR_rest_.

Post-hoc tests showed significantly increased HR_rest_ during the scan from pre to post HIIE (t(19) = 4.71, p < 0.001, d = 1. 054 high pre: 52.03± 6.64 bpm; high post: 58.99 ± 6.95 bpm) and a significantly decreased HR_rest_ from pre to post of the control condition (t(19) = -5.38, p < 0.001, d = -1.20; control pre: 52.49 ± 8.37 bpm; control post: 48.73± 9.01 bpm). No significant HR_rest_ changes were detected from pre to post LIIE (t(19) = 1.65, p = 0.116, d= - 0.37; low pre: 52.44 ± 7.71 bpm; low post: 54.76 ± 9.49 bpm). Comparing the conditions with each other, significant results were found comparing the HIIE versus the control condition (‘high post minus pre’ vs. ‘control post minus pre’; t(19) = 6.85, p < 0.001, d = 1.531; delta high: 6. 96± 6.60 bpm; delta control: -3. 75 ± 3.12 bpm) and comparing the LIIE versus the control condition (‘low post minus pre’ vs. ‘control post minus pre’; t(19) = 4.50, p < 0.001, d = 1.01; delta low: 2.32 ± 6.30 bpm; delta control: -3.75 ± 3.12 bpm). Results were not significant comparing the HIIE versus LIIE condition (‘high post minus pre’ vs. ‘low post minus pre’; t(19) = 2.01, p = 0.059, d = 0.45; delta high: 6.96± 6.60 bpm; delta low: 2.32 ± 6.30 bpm).

#### Exhaustion

Repeated measures ANOVA showed a significant main effect of time (F(1, 19) = 4.61, p = 0.045, η^2^= 0.19), and main effect of condition (F(2, 38) = 3.42, p = 0.043, η^2^= 0.153), but no significant time × condition interaction (F(2, 38) = 1.05, p = 0.361, η^2^= 0. 052) for the variable exhaustion. Post-hoc analysis revealed no significant effects after Bonferroni correction: HIIE condition (t(19) = 2.13, p = 0.046, d = 0.48; high pre: 1.65 ± 1.33; high post: 1.07 ± 1.02); LIIE condition (t(19) = 1.453, p = 0.163, d = 0.32; low pre: 1.02 ± 1.21; low post: 0.62 ± 0.76); control condition (t(19) = 0.322, p = 0.751, d = 0.072; control pre: 1.27 ± 1.22; control post: 1.20 ± 1.13).

### Behavioral Data

#### PANAS

A 2×3 repeated measures ANOVA for PANAS Positive Affect found significant main effects of condition (F(2,38) = 17.29, p < 0.001, η^2^= 0.48) and time (F(1,19) = 28.17, p < 0.001, η^2^ = 0.60), as well as a time x condition interaction (F(2,38) = 17.08, p < 0.001, η^2^ = 0.47). Post hoc comparisons showed significant increases after HIIE (t (19) = 6.42, p < 0.001, Cohen’s d = 1.43; high pre: 30.30 ± 6.67; high post: 38.20 ± 5.21) and after LIIE (t (19) = 5.67, p < 0.001, Cohen’s d = 1.27; low pre: 31.25 ± 6.97; low post: 36.25 ± 6.76). No significant differences were observed from pre to post control conditions. Comparing the conditions with each other, significant results were found when testing HIIE versus the control condition (‘high post minus pre’ vs. ‘control post minus pre’; t (19) = 7.12, p < 0.001, Cohen’s d = 1.59; delta high: 7.90± 5.50; delta control: -0.45 ± 5.73) and testing LIIE versus the control condition (‘low post minus pre’ vs. ‘control post minus pre’ (t (19) = 3.38, p = 0.003, Cohen’s d = 0.756; delta low: 5.0 ± 3.95; delta control: -0.45 ± 5.73). Testing HIIE vs the LIIE condition showed no significant differences (‘high post minus pre’ vs. ‘low post minus pre’; t (19) = 1.89, p = 0.073, Cohen’s d = 0.42; delta high: 7.90± 5.50; delta low: 5.0 ± 3.95) (see **Figure 2A**).

**Figure 2.**
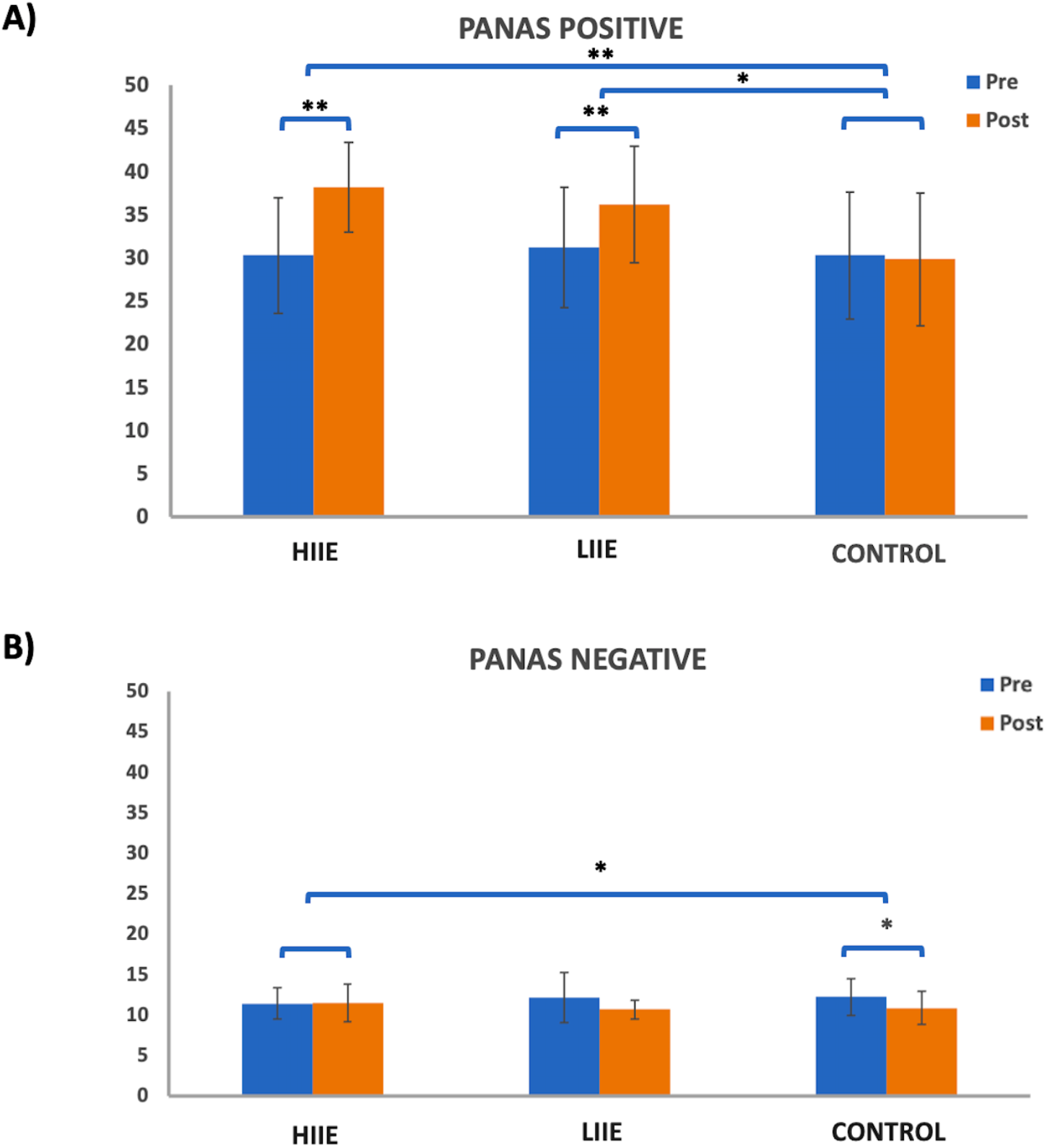
Representation of the PANAS A) Positive and B) Negative scales pre- and post conditions. Whiskers represent the standard deviation. * = p < 0.008 (Bonferroni-corrected); ** = p < 0.001;.

On the other hand, the 2×3 repeated measures ANOVA for the PANAS Negative Affect scale showed a significant main effect of time (F(1,19) = 9.68, p = 0.006, η2 = 0.34) and a significant time x condition interaction (F(2,38) = 5.11, p = 0.011, η2 = 0.21) but no significant main effect of condition (F(2,38) = 0.099, p = 0.906, η2 < 0.01). Post-hoc tests showed significantly decreased PANAS Negative from pre to post control condition (t(19) = -3.94, p = 0.001, d = -0.882; control pre: 12.20 ± 2.62; control post: 10.85 ± 2.03). No significant differences were observed from pre to post of the LIIE condition (t(19) = -2.43, p = 0.025, d = -0.544; low pre: 12.10 ± 3.09; low post: 10.65 ± 1.14) or from pre to post HIIE (t(19) = -0.37, p = 0.716, d= -0.083; high pre: 11.40 ± 1.93; high post: 11.50± 2.33). Comparing the conditions with each other, significant results were found only comparing the high-intensity versus the control condition (‘high post minus pre’ vs. ‘control post minus pre’; t(19) = 3.45, p = 0.003, d = 0.7721; delta high: 0.10± 1.21; delta control: -1.35 ± 1.531). No significant changes were detected when testing the LIIE versus the control condition (‘low post minus pre’ vs. ‘control post minus pre’; t (19) = 0.177, p = 0.862, Cohen’s d = 0.039; delta low: -1.45± 2.67; delta control: -1.35 ± 1.531) or the HIIE vs the LIIE (‘high post minus pre’ vs. ‘low post minus pre’; t (19) = 2.49, p = 0.022, Cohen’s d = 0.56; delta high: 0.10± 1.21; delta low: -1.45± 2.67) (see **Figure 2B**).

#### STAI-State

A 2×3 repeated measures ANOVA for the STAI-State found no significant main effect of condition (F(2,38) = 0.055, p = 0.947, η^2^ < 0.01) and time (F(1,19) = 3.99, p < 0.060, η^2^ = 0.17), as well as no time x condition interaction (F(2,38) = 1.67, p = 0.20, η^2^ = 0.008).

### Voxelwise changes in zfALFF

When comparing the pre-to-post session changes within and between conditions using a 2×3 repeated measures ANOVA, we observed no significant main effects of time (pre versus post), condition (low, high and control), or time × condition interaction effect. However, exploratory post hoc comparisons revealed significant decreases in zfALFF after the HIIE intervention (p<0.05, k (size of the cluster) = 58) in several brain regions including precuneus, orbitofrontal cortex, thalamus, cerebellum, etc. For details see **Figure 3A** and **Table 2A**. Moreover, we found increased (p<0.05, k =86) and decreased (p<0.05, k =68) zfALFF after the HIIE compared to the control condition. The zfALFF increases were found in hypothalamus, periaqueductal gray as well as in the pituitary (see **Figure 3B** and **Table 2B**). Conversely the zfALFF decreases were more widespread and included striatum, precuneus, orbitofrontal cortex, thalamus, and cerebellum (see **Figure 3C** and **Table 2C**). We did not observe any significant time effect within LIIE and control conditions, as well as between conditions (LIIE versus control, and vice versa).

**Figure 3.**
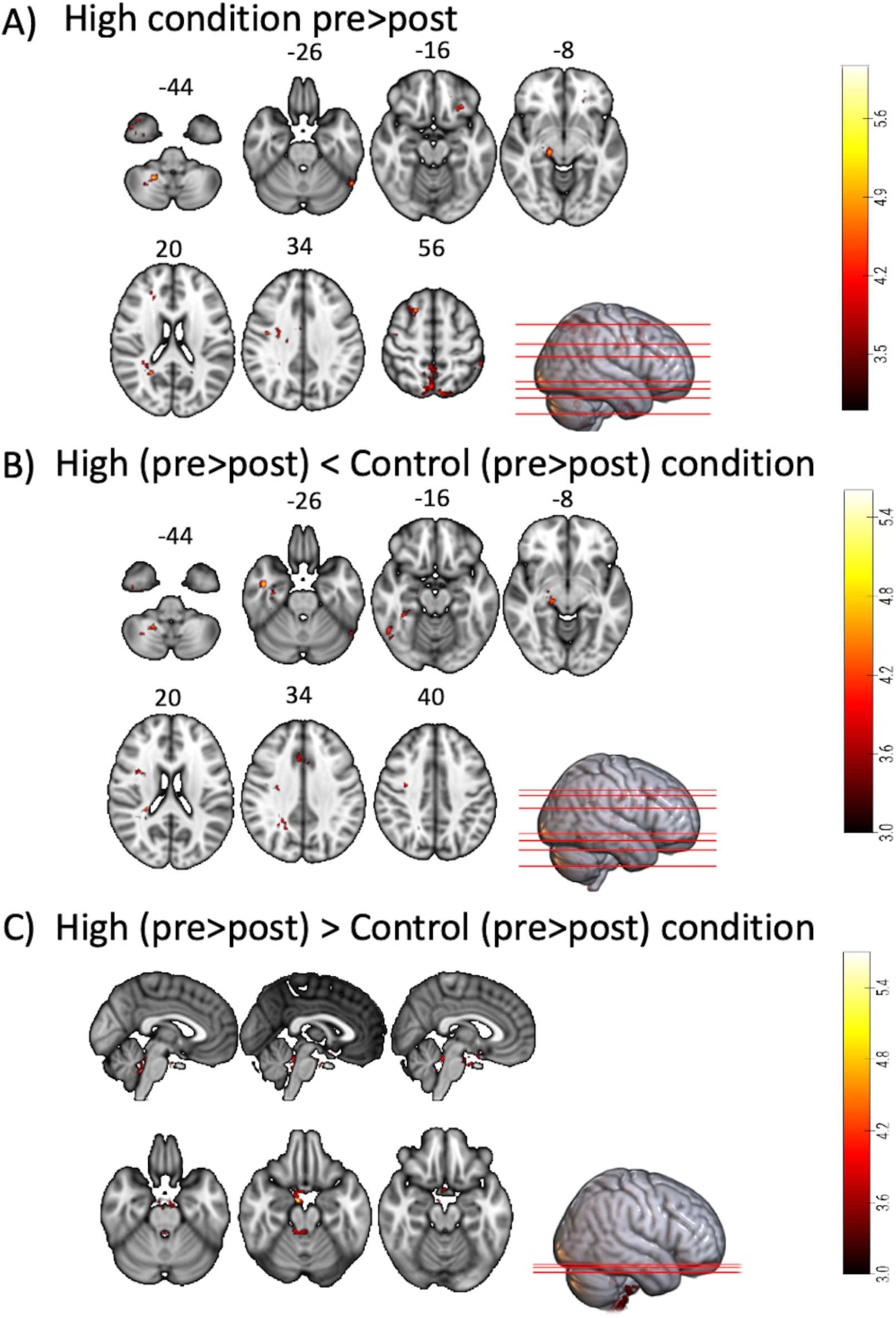
Represents a 2×3 repeatedmeasures ANOVA to compare time and condition effects on zfALFF: A) Results from within condition showing clusters where zfALFF reduced after the HIIE (p<0.05, k=101), B) Results from between condition showing clusters where zfALFF reduced after the HIIE condition compared to control (p<0.05, k=90) and, C) Results from between condition showing clusters where zfALFF increased after the HIIE condition compared to control (p<0.05, k=148). zfALFF = normalized fractional amplitude of low frequency fluctuations; k= cluster size.

**Table 2.** Significant clusters obtained from different posthoc contrasts of 2×3 repeated measures ANOVA.

| No. | Cluster label | Side | Cluster size | MNI coordinates (mm) |  |  | t-value | p-value |
| --- | --- | --- | --- | --- | --- | --- | --- | --- |
|  |  |  | (mm3) | X | Y | Z |  |  |
| A) HIIE Pre > Post |  |  |  |  |  |  |  |  |
| 1 | Cerebellum VIII | R | 195 | 22 | -51 | -45 | 6.09 | <0.001 |
| 2 | Cerebellum IX | R | 101 | 14 | -57 | -33 | 5.61 | 0.025 |
| 3 | Inferior temporal gyrus | L | 120 | -54 | -57 | -24 | 5.43 | 0.009 |
| 4 | Precuneus | L | 117 | -12 | -44 | 15 | 5.36 | 0.011 |
| 5 | Thalamus magnocellular N | R | 134 | 16 | -24 | -8 | 5.34 | 0.004 |
| 6 | Precentral gyrus | R | 355 | 40 | -8 | 33 | 5.19 | <0.001 |
| 7 | Superior frontal gyrus | R | 205 | 21 | 18 | 56 | 5.19 | <0.001 |
| 8 | Precuneus | R | 106 | 24 | -51 | 21 | 5.00 | 0.019 |
| 9 | Middle cingulate gyrus | R | 123 | 10 | -3 | 32 | 4.91 | 0.008 |
| 10 | Inferior orbitofrontal cortex | L | 102 | -21 | 34 | -10 | 4.85 | 0.024 |
| 11 | Superior frontal gyrus | R | 135 | 20 | 33 | 24 | 4.84 | 0.004 |
| 12 | Posterior orbitofrontal cortex | L | 109 | -27 | 26 | -15 | 4.79 | 0.016 |
| 13 | Angular gyrus | R | 143 | 27 | -42 | 32 | 4.70 | 0.003 |
| 14 | Fusiform gyrus | R | 281 | 38 | 3 | -39 | 4.67 | <0.001 |
| 15 | Precuneus | L | 1137 | -10 | -64 | 50 | 4.67 | <0.001 |
| 16 | Inferior parietal gyrus | L | 192 | -52 | -42 | 54 | 4.45 | <0.001 |
| 17 | Supplementary motor area | R | 118 | 4 | 15 | 44 | 4.34 | 0.010 |
| 18 | Superior Parietal gyrus | L | 150 | -8 | -75 | 56 | 4.31 | 0.002 |
| B) HIIE < Control condition |  |  |  |  |  |  |  |  |
| 1 | Middle temporal gyrus | R | 116 | 44 | -3 | -27 | 5.61 | 0.011 |
| 2 | Thalamus ventro-postero-lateral | R | 116 | 16 | -24 | 4 | 5.17 | 0.011 |
| 3 | Cerebellum VIII | R | 145 | 22 | -57 | -40 | 5.09 | 0.002 |
| 4 | Parahippocampal | R | 209 | 36 | -12 | -30 | 4.97 | < 0.001 |
| 5 | Precuneus | R | 186 | 30 | -45 | 26 | 4.64 | <0.001 |
| 6 | Inferior temporal gyrus | R | 135 | 51 | -54 | -15 | 4.63 | 0.004 |
| 7 | Thalamus magnocellular N | R | 122 | 18 | -24 | -8 | 4.59 | 0.008 |
| 8 | Middle cingulate gyrus | R | 125 | 8 | 18 | 32 | 4.56 | 0.007 |
| 9 | Precentral gyrus | R | 138 | 36 | -16 | 30 | 4.53 | 0.004 |
| 10 | Inferior temporal gyrus | L | 108 | -52 | -57 | -24 | 4.45 | 0.017 |
| 11 | Inferior frontal operculum | R | 102 | 32 | 4 | 21 | 4.39 | 0.024 |
| 12 | Fusiform gyrus | R | 119 | 38 | -40 | -15 | 4.26 | 0.010 |
| 13 | Precuneus | R | 104 | 21 | -56 | 33 | 4.17 | 0.021 |

| C) HIIE > Control condition |  |  |  |  |  |  |  |  |
| --- | --- | --- | --- | --- | --- | --- | --- | --- |
| 1 | Hypothalamus | R | 348 | 8 | -6 | -21 | 5.7 | <0.001 |
| 2 | Cerebellum IX | L | 301 | -15 | -38 | -63 | 4.79 | <0.001 |
| 3 | Periaqueductal gray | R | 243 | 10 | -38 | -22 | 4.58 | <0.001 |
| 4 | Parahippocampus | R | 148 | 14 | -20 | -42 | 4.25 | 0.002 |
| 5 | Cerebellum IX | R | 396 | 8 | -45 | -70 | 4.24 | <0.001 |

HIIE = High Intensity Interval Exercise

### Spatial Correlations of fALFF changes with Neurotransmitter Maps

The whole brain analysis of pairwise zfALFF differences of each exercise condition revealed a negative spatial correlation exclusively with the distribution of the endocannabinoid receptor CB1 map after the HIIE condition (Fisher’s z=-0.12, p exact = 0.029, uncorrected; **Figure 4A**). No significant spatial correlation was observed between the pairwise differences in zfALFF and any of the neurotransmitter maps in the LIIE condition or the control condition.

**Figure 4.**
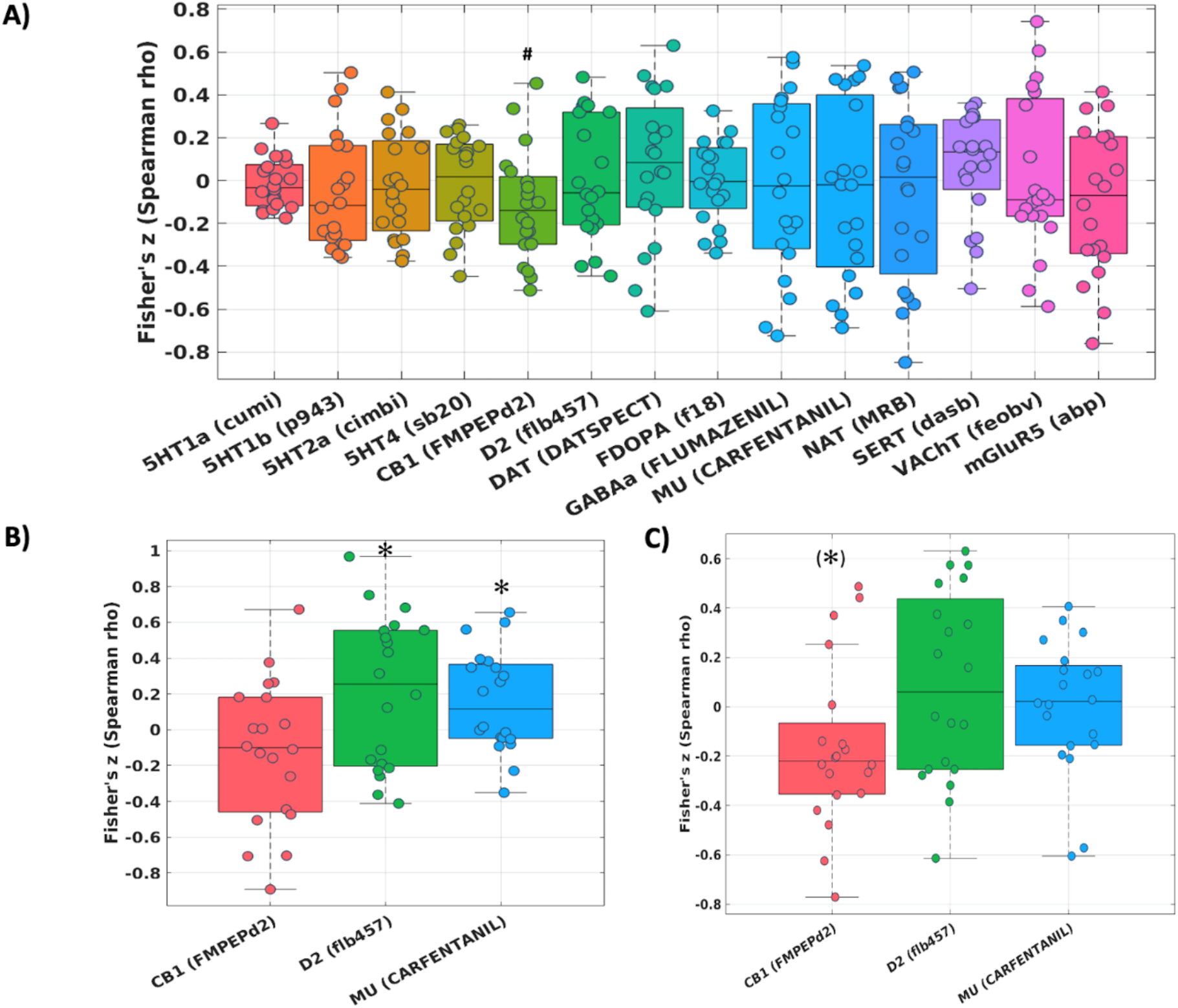
Fisher’s z distribution of the Spearman correlation between the fALFF changes from pre to post in the HIIE condition and neurotransmitters (included in JuSpace toolbox) A) whole brain, B) within the ‘reward network’, and C) within the ‘emotion network’ in trained athletes. * = significant; (*) = trend; # = significant without correction for multiple comparisons.

The hypothesis-driven analysis focusing on the ‘reward network’ showed a significant spatial correlation of the pairwise zfALFF differences and the Mu-opiodergic receptor distribution (Fisher’s z = 0.16, p = 0.027, corrected) after HIIE (**Figure 4B**). Additionally, we observed a significant effect (Fisher’s z = 0.22, p = 0.037, corrected) with the dopamine (D2) receptor distribution after the HIIE condition (**Figure 3B**). Again, no significant correlation was observed within the reward network at the LIIE condition or the control condition.

The hypothesis-driven analysis focusing on the emotion network revealed a trend spatial correlation (Fisher’s z = -0.16, p = 0.064, corrected) between the pairwise zfALFF differences and the CB1 receptor distribution after the HIIE condition (**Figure 4C**).

### fALFF-neurotransmitters Correlations with Mood Changes

We observed a trend positive correlation (r = 0.398; p = 0.082, two-tailed) between the Fishers z values obtained for the association between the zfALFF change and the CB1 receptor distribution within the ‘emotion mask’ and the change in PANAS positive Affect scale (HIIE condition) (**See Supplement Figure S2A**). All other neurotransmitter associations showed no significant or trend correlations with the PANAS Positive or Negative Affect scale changes.

We also observed a trend negative correlation (r = -0.421; p = 0.064, two-tailed) in an explorative analysis (as STAI-State changes were not significant) between the Fishers z values obtained for the association between the zfALFF change and the CB1 receptor distribution within the ‘reward mask’ and the change in STAI-State (HIIE condition) (**See Supplement Figure S2B**). All other neurotransmitter associations showed no significant or trend correlations with the STAI-State changes.

## DISCUSSION

This study examined acute exercise-induced changes in resting-state zfALFF and their spatial cross-correlations with representative PET neurotransmitter distribution maps using the JuSpace toolbox [53] to make indirect inferences about exercise-related acute neurotransmission changes. The within-subject design with three acute training sessions at different activity levels (Control, LIEE, HIIE) in a group of 20 highly trained athletes allowed determining dose-dependent influences of training intensity on resting-state zfALFF changes and their relationship with respective neurotransmitter distributions in the human brain. While both exercise conditions showed the expected positive mood effects (PANAS Positive Affect scale), the HIIE, but not LIIE bouts had post-acute effects on resting-state activation levels in specific brain regions. Associated with these activity changes, results of spatial cross-correlations with representative PET neurotransmitter distribution maps suggest potential involvement of endocannabinoid (via CB1 receptors), dopaminergic (via D2 receptors), and opioidergic (via mu-opioid receptors) neurotransmission after HIIE. In the HIIE condition, trend correlations were found between associations of zfALFF changes with the distribution of CB1 receptors and changes in PANAS Positive Affect and STAI-State, consistent with previous suggestions that endocannabinoid transmission is involved in exercise-induced affective changes.

Exercise had the anticipated mood-improving impact, as evidenced by the higher PANAS Positive Affect following LIIE and HIIE bouts [3, 15, 16]. Meanwhile, no significant increases of PANAS Negative Affect were observed for LIIE and HIIE, indicating that these exercise bouts did not induce post-acute aversive mood states. Indeed, previous evidence suggests that HIIE bouts may induce negative affect during performance (if homeostasis is disrupted to a significant extent) that is followed by a positive affective rebound immediately after exercise [91]. The fact that PANAS Negative Affect only decreased significantly in the control condition is unexpected, given that the participants generally scored in the low range. We can only speculate whether this indicates the athletes’ relieve from passively sitting on the bicycle for an extended time period.

To the best of our knowledge, zfALFF changes have not yet been studied in acute exercise intervention settings, as reported here. Changes in zfALFF changes were exclusively observed after the HIIE condition: Decreases in zfALFF (pre-> post-intervention) were found in precuneus, orbitofrontal cortex, thalamus, and cerebellum. Compared to the control condition, HIIE was associated with differential zfALFF decreases in precuneus, orbitofrontal cortex, thalamus, and cerebellum, whereas differential zfALFF increases were identified in hypothalamus, pituitary, and periaqueductal gray. Comparing these findings with studies using related neuroimaging techniques, we did not replicate previous ASL studies that point towards post-exercise CBF increases in the hippocampus [10] and posterior insula [6]. Yet, the available literature is not consistent, as there were also observations of hippocampal [14, 92] and mesial orbitofrontal CBF decreases [6], the latter being in line with the present result patterns, and also with previous models that suggested transient frontal activity reductions, at least during acute exercise [9]. The significant clusters also show limited overlap with previous rs-fMRI studies which found increased FC in sensorimotor networks [12], or affect-reward, hippocampal, cingulo-opercular, and executive control networks [13], although again, conflicting FC reductions were also observed [14]. Partially consistent with our earlier FC analyses [15, 16], intensity-dependent effects were observed, although in the present case, only the HIIE (not the LIIE) condition produced significant findings. The lack of spatial consistency between our zALFF and previous FC analyses of rs-fMRI data may relate to the fact that the FC methods capture the functional integration between brain regions, and only allow indirect assumptions about the underlying levels. In general, it should be noted that the available neuroimaging studies show substantial methodological variations, e.g. regarding age and fitness level of the participants, duration and intensity of the acute exercise, which may to some degree explain the variable or even contradictory observations. In fact, the relatively intense, but short exercise bouts during HIIE may trigger somewhat different adaptational processes than continuous trainings used in earlier studies.

Interestingly, zfALFF increases were found after HIIE in regions belonging to the hypothalamic-pituitary-adrenal (HPA) axis. One previous study examined hypothalamic CBF, and found no exercise-related effects, but using a continuous, moderate intensity training instead of a HIIE [6]. The hypothalamus integrates signals from other brain nuclei as well as environmental, hormonal, metabolic, and neuronal signals from the periphery [93]. Thereby, vital functions such as energy homeostasis, water balance, and stress are controlled [93], which are essential for adequate responses to exercise, both acutely and chronically [94]. For example, in order to appropriately respond to the rise in body temperature during strenuous exercise, effector responses are triggered by the hypothalamus [95]. Differential zfALFF changes also occurred in the PAG, which is an integral hub region of the opioidergic descending antinociceptive system [96]. Given that pain is a prominent symptom in exercise of high intensity and/or duration it is well conceivable that exercise triggers antinociceptive responses via the PAG [97]. Beyond its role in antinociception, the PAG also fulfils many requirements of a command center for the control of breathing during exercise [98]. It has functional connections to higher brain areas, receives sensory input from contracting muscles, and sends efferent information to brainstem nuclei involved in cardiorespiratory control [98]. In conclusion, the observed zfALFF increases may reflect enhanced regulation of neurohumoral processes, as an adaption to the acute stress during the exercise bouts (see also below).

A particular innovative aspect of the current study is the parallel examination of multiple neurotransmitter effects using rsfMRI after acute exercise bouts with varying intensity. Although the JuSpace approach [53] is *per se* indirect (i.e., spatial cross-correlations of zfALFF changes with representative neurotransmitter maps) and does not allow quantification of endogenous transmitter release (as in PET displacement studies), this MRI-based approach has the unique advantage that several neurotransmitter systems can be tested at the same time in parallel, thereby providing novel insights that help generating hypotheses for future, more transmitter-specific studies. The present results secured the *a priori* proposed roles of the endocannabinoid, dopaminergic, and opioidergic neurotransmitter systems in high-intensity exercise, as suggested by previous research [44, 45, 50]. At whole-brain level, unconstrained exploratory analyses found evidence for pre-post changes exclusively in one of 14 tested neurotransmitter systems, namely the CB1 receptor map, which were moreover restricted to the high-intensity condition. This trend was conserved when limiting analysis to the ‘emotion network’ regions. Follow-up analyses also revealed that the spatial correlation values of the zfALFF changes with the CB1 receptor map were correlated with affective changes in PANAS Positive (but not Negative) Affect, as well as with STAI-State reductions within the the ‘reward network’. In general, these observations are consistent with previous accounts discussing a crucial role of the endocannabinoid system in affective and anxiolytic responses to physical activity [99, 100]. Endocannabinoid receptors are highly expressed in the limbic networks involved in emotional control [47], and animal studies have shown that physical exercise activates the endocannabinoid system, thereby improving mood [50, 100]. Fuss et al. [100] demonstrated in mice that blocking endocannabinoid receptors attenuated the anxiolytic and analgesic effects induced by running, hence, supporting their decisive role in mediating mood changes after acute exercise challenges, whereas this was not the case when the endorphin receptor was blocked. Raichlen et al. [50] found endocannabinoid signaling in healthy athletes to depend on exercise intensity, with peak release occurring at 70-80% of maximum heart rate capacity, which parallels subsequent human studies [49]. Interestingly, the overall spatial correlation between the zfALFF changes and the CB1 map was negative, suggesting a trend for activity decreases across endocannabinoid-rich regions. While CB1 receptors are located presynaptically, and their activation by endocannabinoids generally inhibits the release of other transmitters from these synapses [47], CB1 receptors are expressed on both excitatory and inhibitory synapses which can have complex net effects on signal transmission at system level. There are few human neuroimaging studies that examined the relationship between endocannabinoid modulation and functional brain activity or connectivity. While studies examined acute fMRI effects of phytocannabinoid drugs, especially Δ9-trans-Tetrahydrocannabinol (THC) [101], their global, unselective activation of CB1 receptors does not mimic natural physiological processes. Some fMRI studies examined the impact of genetic polymorphisms that control the production of fatty acid amide hydrolase (FAAH), the main enzyme for AEA degredation. Similar to FAAH knockout mice, individuals with a missense mutation in the FAAH C385A gene show reduced FAAH protein expression and enzymatic activity which results in increased AEA levels in the brain, and is linked with greater fear extinction, enhanced fronto-limbic connectivity and decreased anxiety-like behaviors in both mouse and humans [102]. These individuals also show dampened amygdala responses to threatening faces (along with increased ventral striatal responses to rewards: [103]). Moreover, there are initial observations that amygdala responses to threatening faces are negatively correlated with the availability of FAAH in the medial prefrontal cortex, cingulate and hippocampus, as measured with a recent PET radiotracer [104]. These findings may support a potential dampening effect of endocannabinoid release, especially in response to aversive stimuli.

The analysis restricted to the ‘reward network’ also revealed significant dopaminergic and μ-opiodergic neurotransmission effects after the high-intensity exercise condition, confirming our *a priori* hypotheses. On the one hand, results showed a positive association, suggesting higher brain activity (as indicated by zfALFF), in the D2-rich regions of the reward network. This positive correlation parallels findings from the original JuSpace study, which validated the toolbox with data from an acute risperidone challenge study [53]. If these brain regions show a higher level of baseline activity immediately after exercise bouts, this may help to explain why earlier studies observed blunted ventral striatal responses during monetary reward anticipation and receipt after treadmill exercise [4]. Meanwhile, specific insights into exercise-induced dopamine release changes within motivation-related regions will need future PET displacement studies. On the other hand, there was also a positive spatial correlation between the zfALFF and the mu-opioid distribution map within the reward network regions. Both neurotransmitter systems are assumed to play complementary roles in reward processing, with dopamine being more closely related to motivational-energetic (‘wanting’) aspects, and opioids being more closely related to hedonic-consummatory aspects (‘liking’) of reward [105]. Indeed, joint opioidergic and dopaminergic neurotransmission was detected in the ventral striatum of rats: A rostro-dorsally located ‘opioid hedonic hotspot’ in the nucleus accumbens, mediating behavioral ‘liking’ reactions via endorphinergic transmission [106]. This ‘liking’ hotspot could be distinguished from a separate, more caudally located hotspot for ‘wanting’ behavior mediated via dopaminergic transmission [106]. Interestingly, one human study observed that the individual strength of opioid release after moderate exercise training correlated positively with the brain response of reward network areas (including ventral striatum) while viewing palatable, as compared to nonpalatable food pictures, which would be consistent with an opioidergic mechanisms to hedonic aspects of reward processing [46]. Although we do not have behavioral results to distinguish the representations of ‘liking’ and ‘wanting’ in the ventral striatum, it is noteworthy that our analyses indicate a common representation of these two neurotransmitter systems and an effect of HIIE on both. This finding is also consistent with observations that central opioid release appears to be dependent on the intensity of exercise [44, 45]. Previous work from our group identified exercise-induced opioidergic transmitter release, as measured with 6-O-(2-[18F]fluoroethyl)-6-O-desmethyldiprenor-phine ([18F]DPN) PET [22]. Importantly, [18F]DPN binding changes correlated with VAS euphoria change scores [22]. Why such a relationship was not found with the current indirect analysis approach remains a matter of speculation and should be addressed in future studies. Another study reported that increased opioid release was correlated with increased euphoria after a moderate intensity continuous exercise, but increased negative affect after high-intensity interval training [45], which may indicate that at very high levels of exercise strain the stress-reducing effects of opioidergic neurotransmission become more important. In this context it is interesting to revisit an abovementioned observation from the present study: It is intriguing that zfALFF changes were found in the hypothalamus and the PAG, both of which are known as core regions of the central opioidergic neurotransmitter system, but not covered by the JuSpace maps. The hypothalamus is the region where β-endorphins are produced and released, in particular in the pro-opio-melano-cortin (POMC) neurons located in the arcuate hypothalamic nucleus (ARH) and, to a lesser extent, in the nucleus of the solitary tract [107]. β-endorphins from the hypothalamic POMC neurons in the ARH are released inside the central nervous system, notably to limbic structures, hypothalamic and thalamic sites, and several brainstem nuclei [107]. On the other hand, β-endorphins produced in the nucleus solitarius project axons to the spinal cord. In addition to these central acting β-endorphins, those from the pituitary are released into the peripheral systemic circulation [107]. Activation of the hypothalamic-pituitary-adrenal (HPA) axis by physical exercise has been demonstrated by increased endorphin levels in the pituitary [108, 109] and elevated plasma immunoreactive beta-endorphin/beta-lipotropin levels [110, 111]. Differential zfALFF changes in the PAG are equally interesting as the region is a crucial hub region of the descending antinociceptive system which is opoidergically mediated and modulated by exercise, as shown by our group in human athletes after strenuous exercise [97]. In line with our observations, it was demonstrated in mice after exposure to forced walking stress that β-endorphin levels are increased in PAG and/or medial basal hypothalamus and may be involved in stress-induced analgesia [112]. Meanwhile, extensive expression of CB1 and CB2 receptors of the endocannabinoid system has been found in limbic regions and also in the hypothalamus [113]. Furthermore, CB1 receptor signaling has been shown to have an effect on the activation of the HPA axis [114]. One thus needs to consider that there are mutual interactions between the endocannabinoid systems and the opioidergic [38, 115] and dopaminergic [115] systems in the human brain, with indications that the opioid system is involved in the increase of AEA and OEA following exercise [38]. Yet, with the currently available technical approaches, these complex interrelationships are difficult to disentangle in humans.

The additional observation that changes of spatial correlation between CB1 and zfALFF map within emotion-and reward-related networks showed trend correlations with increases in positive mood and reductions in anxiety, suggests that functional interpretation should be made with caution: Given that the group-level analyses found a general negative spatial correlation between zfALFF and CB1 maps, this would imply that participants with stronger positive mood increases / anxiety decreases after the HIIE showed *less* pronounced zfALFF *decreases* in endocannabinoid-rich brain networks, i.e. by inference, less dampening of brain activity. One may speculate that these participants experienced less need for downregulation of brain activity due to a lower exercise burden, allowing for a stronger affective rebound. Notably, most evidence for an endocannabinoid modulation of affect originates from animal studies that examined aversive situations, i.e. the anxiolytic or anti-stress role of endocannabinoid signaling [47]. We did not find exercise-induced reduction in post-exercise negative affect, but this was difficult because the participants already performed at floor level for the PANAS Negative Affect scale. Possibly, objective measurements of physiological responses to affective stimuli (like task paradigms in the abovementioned FAAH-related fMRI studies) can help to detect more subtle changes beyond subjective experience. Most importantly, direct information about exercise-induced endocannabinoid activation (and its inter-and intraindividual variation) is needed. While PET imaging of the endocannabinoid system is still evolving, and has to deal with methodological and regulatory limitations, future studies investigating the affective influence of acute exercise bouts should at least include peripheral measures of endocannabinoid release [51].

This study is not without limitations: The JuSpace method is by definition indirect and will only provide indirect indications about the involvement of neurotransmitter systems involved in exercise challenges. We constrained our analyses of the ‘emotion network’ and the ‘reward network’ to three transmitter systems described in the context of affect and reward processing. The reported effects were only detectable in the high-intensity condition and effect sizes were too weak to survive corrections for multiple testing of all available PET transmitter maps. Therefore, future studies of this kind should attempt to measure larger samples in order to detect further neurotransmitter involvement related to exercise. We can only speculate on the association with neurotransmitter effects in those regions like the hypothalamus and the PAG which were not covered by the PET tracer distribution maps. Finally, future studies should be enrolled in larger samples including all sexes, which was not the case in this pilot examination.

In summary, this is the first attempt to study exercise-induced neurotransmission non-invasively and in a rather holistic manner using rs-fMRI in humans. Our approach of studying zfALFF changes in the context of acute exercise bouts differing in intensity has provided novel information on the neurobiology of exercise, extending previous evidence on the particular relevance of the opioidergic, the dopaminergic and the endocannabinoid systems in high-intensity physical activity. In exploratory statistical analyses, there was a link between changes in mood and the endocannabinoid system, consistent with its role as an affective modulator system, but further research is needed to substantiate this claim in humans. Interactions between these transmitter systems have been described and should be studied further *in vivo*, e.g. between endocannabinoid signaling and endorphin [115, 116] or dopamine release [115, 117] in the specific context of exercise physiology.

## Data availability statement

Data can only be made available after contacting participating volunteers and obtaining their consent to submit data in anonymized form (as per ethics’ approval). Depending on the decision of the volunteers, this may result in smaller samples.

## Abbreviations

BOLD: Blood oxygenation level-dependent
MRI: Magnetic resonance imaging
RsfMRI: Resting state functional MRI
PET: Positron emission tomography
fALFF: fractional amplitude of low frequency fluctuations
PANAS: Positive and negative affect scale
FDR: False discovery rate
PE: Physical exercise
HR: Heart rate
HR_int_: Heart rate during exercise intervention
HR_rest_: Heart rate during resting state functional MRI
PE: Physical exercise
RER: Respiratory exchange ratio
ROI: Region of interest
VO_2_: Oxygen uptake
VO_2max_: Maximal oxygen uptake (mL/min/kg)

