## Supplementary Material for "Fractional Amplitude of Low-Frequency Fluctuations Associated with Endocannabinoid, μ-Opioid and Dopamine Receptor Distributions in the Central Nervous System after High-Intensity Exercise Bouts"

**SUPPLEMENT**

Detailed Description of the fMRIPrep Pipeline (Boilerplate)

Preprocessing of structural and functional MRI data was performed using fMRIPrep 20.2.6 (Esteban, Markiewicz, et al. (2018); Esteban, Blair, et al. (2018); RRID:SCR_016216), which is based on Nipype 1.7.0 (Gorgolewski et al. (2011); Gorgolewski et al. (2018); RRID:SCR_002502).

Anatomical data preprocessing

A total of 6 T1-weighted (T1w) images were found within the input BIDS dataset. All of them were corrected for intensity non-uniformity (INU) with N4BiasFieldCorrection (Tustison et al. 2010), distributed with ANTs 2.3.3 (Avants et al. 2008, RRID:SCR_004757). The T1w-reference was then skull-stripped with a Nipype implementation of the antsBrainExtraction.sh workflow (from ANTs), using OASIS30ANTs as target template. Brain tissue segmentation of cerebrospinal fluid (CSF), white-matter (WM) and gray-matter (GM) was performed on the brain-extracted T1w using fast (FSL 5.0.9, RRID:SCR_002823, Zhang, Brady, and Smith 2001). A T1w-reference map was computed after registration of 6 T1w images (after INU-correction) using mri_robust_template (FreeSurfer 6.0.1, Reuter, Rosas, and Fischl 2010). Brain surfaces were reconstructed using recon-all (FreeSurfer 6.0.1, RRID:SCR_001847, Dale, Fischl, and Sereno 1999), and the brain mask estimated previously was refined with a custom variation of the method to reconcile ANTs-derived and FreeSurfer-derived segmentations of the cortical gray-matter of Mindboggle (RRID:SCR_002438, Klein et al. 2017). Volume-based spatial normalization to two standard spaces (MNI152NLin2009cAsym, MNI152NLin6Asym) was performed through nonlinear registration with antsRegistration (ANTs 2.3.3), using brain-extracted versions of both T1w reference and the T1w template. The following templates were selected for spatial normalization: ICBM 152 Nonlinear Asymmetrical template version 2009c [Fonov et al. (2009), RRID:SCR_008796; TemplateFlow ID: MNI152NLin2009cAsym], FSL's MNI ICBM 152 non-linear 6th Generation Asymmetric Average Brain Stereotaxic Registration Model [Evans et al. (2012), RRID:SCR_002823; TemplateFlow ID: MNI152NLin6Asym].

Functional data preprocessing

For each of the 6 BOLD runs found per subject (across all tasks and sessions), the following preprocessing was performed. First, a reference volume and its skull-stripped version were generated using a custom methodology of fMRIPrep. A B0-nonuniformity map (or fieldmap) was directly measured with an MRI scheme designed with that purpose (typically, a spiral pulse sequence). The fieldmap was then co-registered to the target EPI (echo-planar imaging) reference run and converted to a displacements field map (amenable to registration tools such as ANTs) with FSL's fugue and other SDCflows tools. Based on the estimated susceptibility distortion, a corrected EPI (echo-planar imaging) reference was calculated for a more accurate co-registration with the anatomical reference. The BOLD reference was then co-registered to the T1w reference using bbregister (FreeSurfer) which implements boundary-based registration (Greve and Fischl 2009). Co-registration was configured with six degrees of freedom. Head-motion parameters with respect to the BOLD reference (transformation matrices, and six corresponding rotation and translation parameters) are estimated before any spatiotemporal filtering using mcflirt (FSL 5.0.9, Jenkinson et al. 2002). The BOLD time-series (including slice-timing correction when applied) were resampled onto their original, native space by applying a single, composite transform to correct for head-motion and susceptibility distortions. These resampled BOLD time-series will be referred to as preprocessed BOLD in original space, or just preprocessed BOLD. The BOLD time-series were resampled into standard space, generating a preprocessed BOLD run in MNI152NLin2009cAsym space. First, a reference volume and its skull-stripped version were generated using a custom methodology of fMRIPrep. Automatic removal of motion artifacts using independent component analysis (ICA-AROMA, Pruim et al. 2015) was performed on the preprocessed BOLD on MNI space time-series after removal of non-steady state volumes and spatial smoothing with an isotropic, Gaussian kernel of 6mm FWHM (full-width half-maximum). Corresponding "non-aggresively" denoised runs were produced after such smoothing. Additionally, the "aggressive" noise-regressors were collected and placed in the corresponding confounds file. Several confounding time-series were calculated based on the preprocessed BOLD: framewise displacement (FD), DVARS and three region-wise global signals. FD was computed using two formulations following Power (absolute sum of relative motions, Power et al. (2014)) and Jenkinson (relative root mean square displacement between affines, Jenkinson et al. (2002)). FD and DVARS are calculated for each functional run, both using their implementations in Nipype (following the definitions by Power et al. 2014). The three global signals are extracted within the CSF, the WM, and the whole-brain masks. Additionally, a set of physiological regressors were extracted to allow for component-based noise correction (CompCor, Behzadi et al. 2007). Principal components are estimated after high-pass filtering the preprocessed BOLD time-series (using a discrete cosine filter with 128s cut-off) for the two CompCor variants: temporal (tCompCor) and anatomical (aCompCor). tCompCor components are then calculated from the top 2% variable voxels within the brain mask. For aCompCor, three probabilistic masks (CSF, WM and combined CSF+WM) are generated in anatomical space. The implementation differs from that of Behzadi et al. in that instead of eroding the masks by 2 pixels on BOLD space, the aCompCor masks are subtracted a mask of pixels that likely contain a volume fraction of GM. This mask is obtained by dilating a GM mask extracted from the FreeSurfer's aseg segmentation, and it ensures components are not extracted from voxels containing a minimal fraction of GM. Finally, these masks are resampled into BOLD space and binarized by thresholding at 0.99 (as in the original implementation). Components are also calculated separately within the WM and CSF masks. For each CompCor decomposition, the k components with the largest singular values are retained, such that the retained components' time series are sufficient to explain 50 percent of variance across the nuisance mask (CSF, WM, combined, or temporal). The remaining components are dropped from consideration. The head-motion estimates calculated in the correction step were also placed within the corresponding confounds file. The confound time series derived from head motion estimates and global signals were expanded with the inclusion of temporal derivatives and quadratic terms for each (Satterthwaite et al. 2013). Frames that exceeded a threshold of 0.5 mm FD or 1.5 standardised DVARS were annotated as motion outliers. All resamplings can be performed with a single interpolation step by composing all the pertinent transformations (i.e. head-motion transform matrices, susceptibility distortion correction when available, and co-registrations to anatomical and

output spaces). Gridded (volumetric) resamplings were performed using antsApplyTransforms (ANTs), configured with Lanczos interpolation to minimize the smoothing effects of other kernels (Lanczos 1964). Non-gridded (surface) resamplings were performed using mri_vol2surf (FreeSurfer).
Many internal operations of fMRIPrep use Nilearn 0.6.2 (Abraham et al. 2014, RRID:SCR_001362), mostly within the functional processing workflow. For more details of the pipeline, see the section corresponding to workflows in fMRIPrep's documentation.

Copyright Waiver

The above boilerplate text was automatically generated by fMRIPrep with the express intention that users should copy and paste this text into their manuscripts unchanged. It is released under the CC0 license.

<u>Physiological Data Analysis</u>

| **$HR_{int}$ [bpm]** | | **High (Mean ± SD)** | **Low (Mean ± SD)** | **p-value** | **T-value** | **df** | **Cohens d** |
|---|---|---|---|---|---|---|---|
| Interval 1 | load | 167 ± 9 | 138 ± 10 | <0.001 | 13.261 | 19 | 2.965 |
| | recovery | 139 ± 10 | 136 ± 9 | 0.669 | 1.436 | 19 | 0.321 |
| Interval 2 | load | 176 ± 7 | 144 ± 11 | <0.001 | 11.817 | 19 | 2.642 |
| | recovery | 145 ± 9 | 140 ± 11 | 0.641 | 1.461 | 19 | 0.327 |
| Interval 3 | load | 180 ± 7 | 146 ± 10 | <0.001 | 13.693 | 19 | 3.062 |
| | recovery | 151 ± 11 | 142 ± 10 | 0.028 | 3.029 | 19 | 0.677 |
| Interval 4 | load | 184 ± 6 | 148 ± 11 | <0.001 | 13.917 | 19 | 3.112 |
| | recovery | 151 ± 11 | 143 ± 11 | 0.088 | 2.493 | 19 | 0.557 |

***Table S1.*** *Results of statistical analysis of $HR_{int}$ data during exercise. Paired t-tests with Bonferroni correction were performed. p-value = Bonferroni-corrected p-value.*

| La [mmol/L] | | High (Mean ± SD) | Low (Mean ± SD) | p-value | T-value | df | Cohens d |
|---|---|---|---|---|---|---|---|
| Interval 1 | load | 5.7 ± 1.2 | 1.5 ± 0.6 | <0.001 | 20.457 | 19 | 4.574 |
| | recovery | 5.4 ± 1.6 | 1.4 ± 0.7 | <0.001 | 13.754 | 18 | 3.155 |
| Interval 2 | load | 7.2 ± 1.9 | 1.5 ± 0.7 | <0.001 | 16.816 | 19 | 3.760 |
| | recovery | 6.5 ± 2.4 | 1.4 ± 0.6 | <0.001 | 10.774 | 19 | 2.409 |
| Interval 3 | load | 8.0 ± 2.4 | 1.4 ± 0.6 | <0.001 | 14.126 | 19 | 3.159 |
| | recovery | 7.3 ± 2.7 | 1.4 ± 0.6 | <0.001 | 10.189 | 18 | 2.338 |
| Interval 4 | load | 8.6 ± 2.8 | 1.5 ± 0.5 | <0.001 | 11.963 | 18 | 2.745 |
| | recovery | 8.1 ± 3.2 | 1.4 ± 0.6 | <0.001 | 10.363 | 19 | 2.317 |

***Table S2.*** *Results of statistical analysis of lactate data during exercise. Paired t-tests with Bonferroni correction were performed. p-value = Bonferroni-corrected p-value.*

| RPE | | High (Mean ± SD) | Low (Mean ± SD) | p-value | T-value | df | Cohens d |
|---|---|---|---|---|---|---|---|
| Interval 1 | load | 15 ± 1 | 11 ± 2 | <0.001 | 7.535 | 19 | 1.685 |
| | recovery | 10 ± 1 | 11 ± 2 | 1.000 | -0.396 | 19 | -0.089 |
| Interval 2 | load | 16 ± 1 | 12 ± 1 | <0.001 | 9.469 | 19 | 2.117 |
| | recovery | 11 ± 2 | 11 ± 2 | 1.000 | 0.954 | 19 | 0.213 |
| Interval 3 | load | 16 ± 1 | 12 ± 1 | <0.001 | 8.976 | 19 | 2.007 |
| | recovery | 12 ± 2 | 11 ± 2 | 0.057 | 2.698 | 19 | 0.603 |
| Interval 4 | load | 17 ± 1 | 12 ± 2 | <0.001 | 11.786 | 19 | 2.636 |
| | recovery | 12 ± 2 | 11 ± 2 | 1.000 | 0.879 | 19 | 0.197 |

***Table S3.*** *Results of statistical analysis of RPE data during exercise. Paired t-tests with Bonferroni correction were performed. p-value = Bonferroni-corrected p-value.*

| Subject | First Rise [W] | $D_{max}$ [W] | High | | | | Low | | | |
|---|---|---|---|---|---|---|---|---|---|---|
| | | | Target_Load 110% $D_{max}$ [W] 4*4 min | Load (Mean ± SD) | Target_Recovery 60% $D_{max}$ [W] 4*3 min | Recovery (Mean ± SD) | Target_Load 100% First Rise [W] 4*4 min | Load (Mean ± SD) | Target_Recovery 90% First Rise [W] 4*3 min | Recovery (Mean ± SD) |
| 01 | 160 | 231 | 254 | 254 ± 5 | 139 | 139 ± 5 | 160 | 160 ± 0 | 144 | 144 ± 0 |
| 02 | 180 | 237 | 261 | 261 ± 3 | 142 | 142 ± 0 | 180 | 180 ± 0 | 162 | 162 ± 0 |
| 03 | 220 | 299 | 329 | 329 ± 5 | 179 | 179 ± 7 | 220 | 220 ± 0 | 198 | 198 ± 0 |
| 04 | 180 | 257 | 283 | 283 ± 0 | 154 | 154 ± 0 | 180 | 180 ± 1 | 162 | 162 ± 0 |
| 05 | 200 | 270 | 298 | 298 ± 3 | 162 | 162 ± 4 | 200 | 200 ± 2 | 180 | 180 ± 1 |
| 06 | 220 | 284 | 313 | 313 ± 0 | 170 | 170 ± 0 | 220 | 220 ± 1 | 198 | 198 ± 1 |
| 07* | 280 | 340 | 374 | - | - | - | 280 | 280 ± 1 | 252 | 252 ± 1 |
| 08 | 180 | 246 | 270 | 270 ± 4 | 147 | 147 ± 3 | 180 | 180 ± 2 | 162 | 162 ± 0 |
| 09 | 240 | 288 | 317 | 317 ± 3 | 173 | 173 ± 4 | 240 | 240 ± 3 | 216 | 216 ± 1 |
| 10 | 220 | 280 | 307 | 307 ± 3 | 168 | 168 ± 5 | 220 | 220 ± 1 | 198 | 198 ± 1 |
| 11 | 200 | 285 | 313 | 313 ± 4 | 171 | 171 ± 4 | 200 | 200 ± 0 | 180 | 180 ± 1 |
| 12 | 300 | 345 | 379 | 379 ± 0 | 207 | 207 ± 5 | 300 | 300 ± 1 | 270 | 270 ± 1 |
| 13 | 220 | 280 | 308 | 308 ± 0 | 168 | 168 ± 4 | 220 | 220 ± 1 | 198 | 198 ± 1 |

| | | | | | | | | | | |
|---|---|---|---|---|---|---|---|---|---|---|
| 14 | 140 | 222 | 244 | 244 ± 0 | 133 | 133 ± 3 | 140 | 140 ± 0 | 126 | 126 ± 0 |
| 15 | 180 | 233 | 256 | 256 ± 4 | 140 | 140 ± 3 | 180 | 180 ± 0 | 162 | 162 ± 0 |
| 16 | 240 | 297 | 327 | 327 ± 0 | 178 | 178 ± 0 | 240 | 240 ± 3 | 216 | 216 ± 1 |
| 17 | 200 | 337 | 371 | 371 ± 4 | 202 | 201 ± 12 | 200 | 200 ± 2 | 180 | 180 ± 1 |
| 18 | 260 | 330 | 363 | 363 ± 8 | 198 | 198 ± 6 | 260 | 260 ± 1 | 234 | 234 ± 1 |
| 19 | 180 | 232 | 255 | 255 ± 4 | 139 | 139 ± 5 | 180 | 180 ± 0 | 162 | 162 ± 0 |
| 20 | 180 | 240 | 264 | 264 ± 3 | 144 | 144 ± 3 | 180 | 180 ± 1 | 162 | 162 ± 0 |

***Table S4.*** *Individual power value in W for First Rise and $D_{max}$, determined in performance diagnostics. Target_Load and Target_Recovery values are showing the calculated target values for the corresponding intervals during the high-intensity and low-intensity interventions. The values in the respective following column represent the mean ± standard deviation of the actual power values during the intervention. * Missing values due to technical problems exporting the performance data after the intervention.*

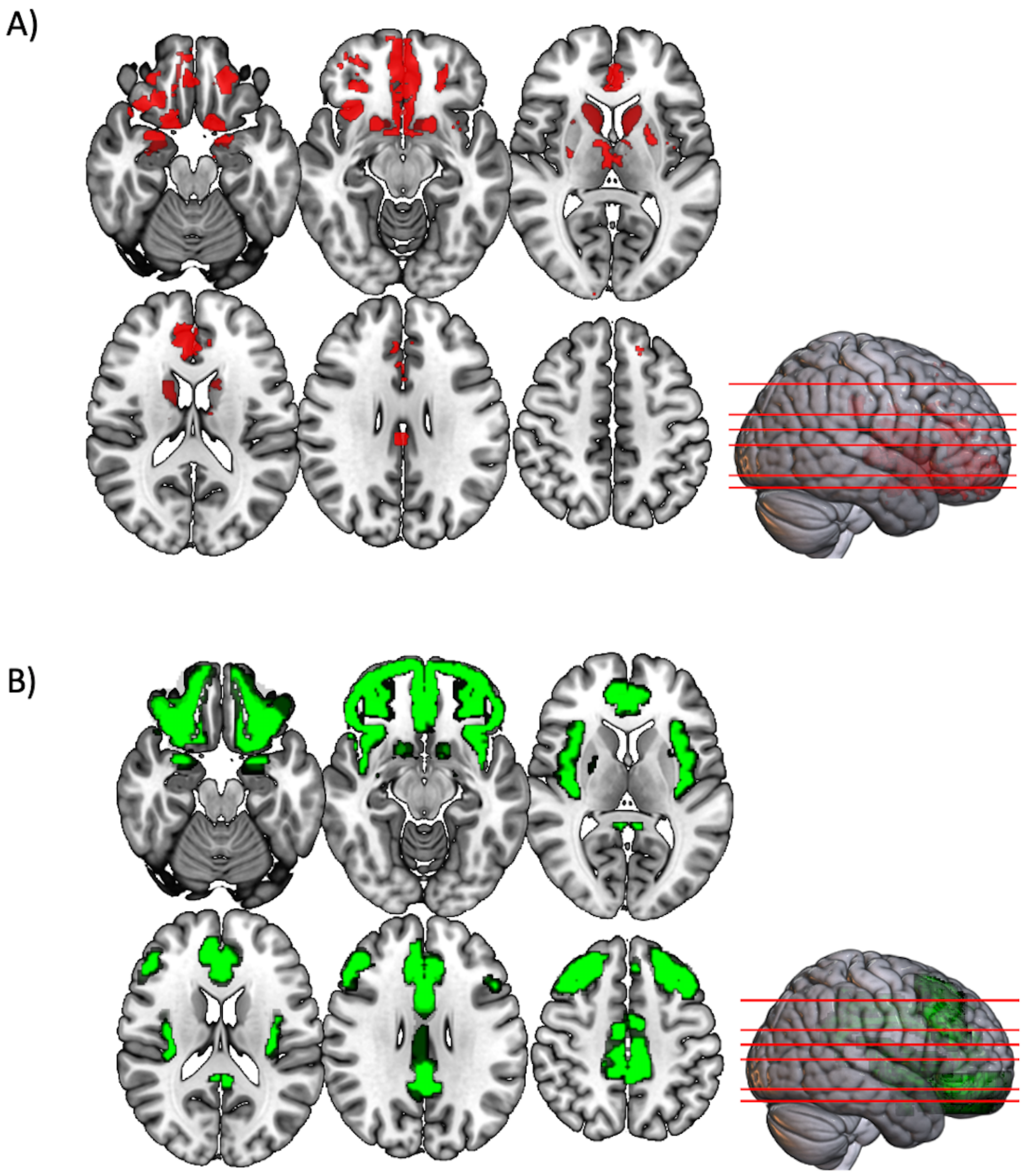


**Figure S1: Reward and Emotion Network Masks.** Representation of masks related to A) reward network (red) and B) emotion network (green).

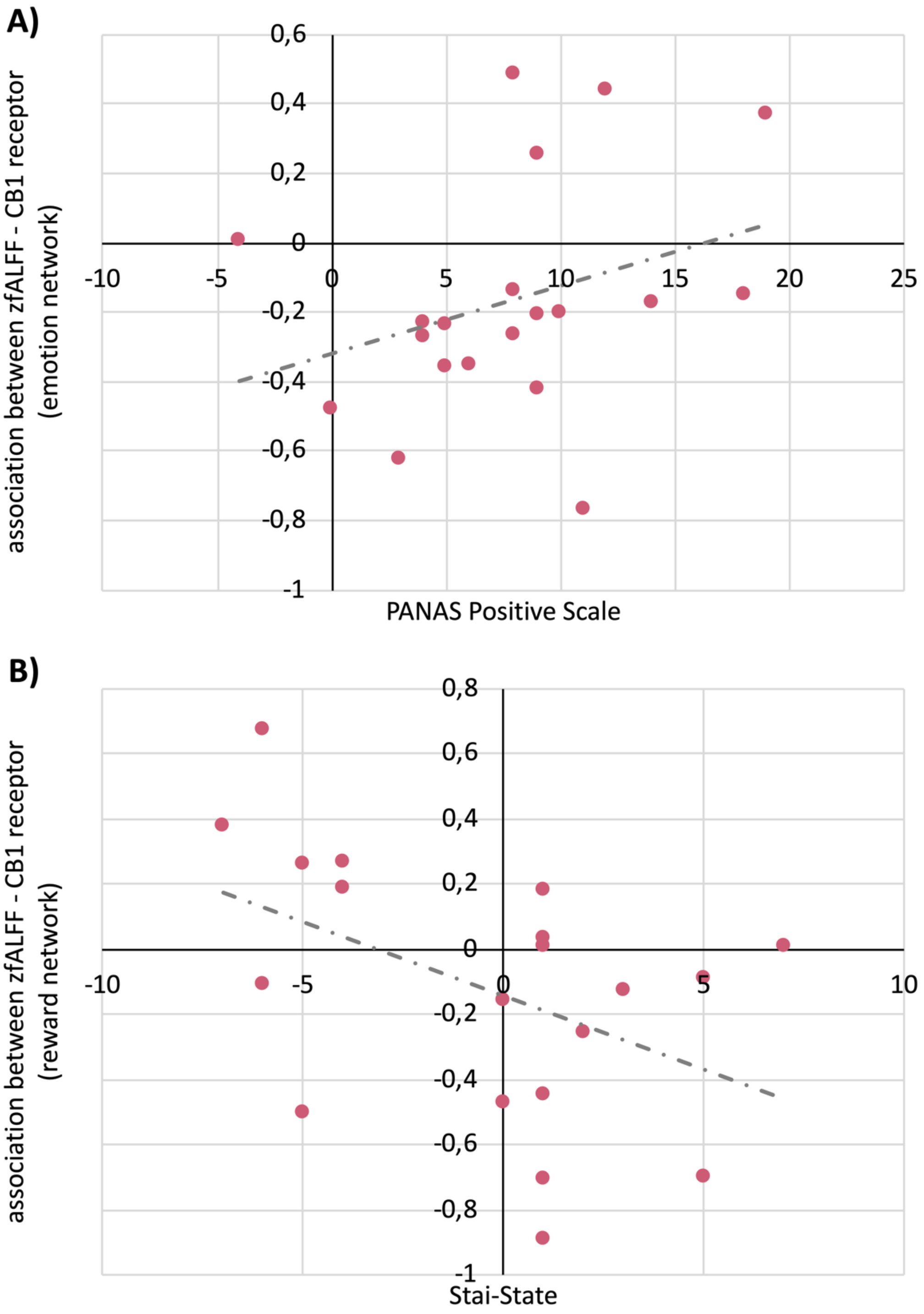


**Figure S2: fALFF-neurotransmitters Correlations with Mood Changes.** A) Trend positive correlation between the Fishers z values obtained for the association between the zfALFF change and the CB1 receptor distribution within the 'emotion mask' and the change in PANAS positive Affect scale (HIIE condition). B) Trend negative correlation between the Fishers z values obtained for the association between the zfALFF change and the CB1 receptor distribution within the 'reward mask' and the change in STAI-State (HIIE condition).
